# Single-Sequence, Structure Free Allosteric Residue Prediction with Protein Language Models

**DOI:** 10.1101/2024.10.03.616547

**Authors:** Gokul R. Kannan, Brian L. Hie, Peter S. Kim

## Abstract

Large language models trained on protein amino acid sequences have shown the ability to learn general coevolutionary relationships at scale, which in turn contain useful structural and functional information. Here we show that attention maps, matrices of learned pairwise relationships between residues, also include information about allostery. This enables prediction of allosteric relationships with no task-specific training, requiring only a single input sequence and no structural information. Attention maps outperform state-of-the-art structure-based and sequence coevolution-based allosteric residue prediction models on a well-curated benchmark set of 24 allosteric proteins. For K-Ras, an allosterically regulated GTPase, attention maps correlate best with allosteric residues influencing binding identified in deep mutational scanning data. For the beta-2 adrenergic receptor, an allosterically regulated GPCR, attention maps correlate best when compared to experimental alanine-scanning mutational data identifying allosteric relationships influencing signaling. These results enable allosteric relationship prediction in a single-sequence, structure-free manner.

## Main

The evolutionary trajectory of a protein sequence reflects its optimization along multiple axes, including structure, function, and stability. Recent evidence suggests that protein language models learn general coevolutionary relationships from training on sequence information alone, enabling these models to infer three-dimensional structural information, predict evolutionary trajectories over both short and long timescales, and navigate complex protein fitness landscapes.^1–7^

Remarkably, these abilities emerge from a simple masked language modeling training objective. This involves masking a single amino acid residue within a protein sequence and asking the model to predict the identity of the masked residue. By training on millions of sequences using models with billions of parameters, these models develop an understanding of contextual relationships between amino acids. Language models employ self-attention mechanisms to identify these relationships, including interactions involving distant parts of a sequence.^8^ Self-attention allows the model to weigh the importance of each part of the input sequence differently when predicting a particular token (an amino acid in a protein language model). These pairwise relationships between every pair of tokens in the sequence are learned during training as an *L x L* attention map matrix, where L is the sequence length. Most modern language models are implemented with the Transformer architecture, which consists of sequential neural network layers where each layer computes the attention operation in parallel on the same input (referred to a multiple “heads” of attention).^8^

Attention maps are useful representations of biochemical phenomena. These learned pairwise relationships have been shown to extract structural information about proteins, which is possible because residues in contact are more likely to coevolve with each other to maintain structural coherence.^1,6,9,10^ Thus, coevolution of residue pairs often translates to high attention values between those pairs, which in turn can be used to extract contact map predictions.

In this study, we sought to characterize residue pairs that have high values in the attention maps but are distant in structure. In particular, we hypothesized that these could represent allosteric information. Allostery, termed by Monod as the ‘second secret of life’ after the genetic code, has evolved in many proteins whereby changes such as covalent modifications, binding of ligands, or amino acid mutations can modify a protein’s function despite these changes occurring outside the protein’s active site.^11^ In the most general sense, allostery occurs when a modification at one site (the “allosteric site”) induces a change at some non-overlapping coupled site (the “active site”).^11^ In the classic example, allostery in hemoglobin allows it to maximally load and unload oxygen in oxygen-rich and oxygen-deficient environments respectively. Allostery has been exploited in therapeutic discovery extensively by drugs such as benzodiazepines (GABA modulators) and sotorasib (a KRAS G12C modulator). While important in drug discovery and prevalent in signaling systems, discovery of new allosteric sites and relationships remains difficult even for very well-studied proteins.

Allosteric regulation appears to be captured in coevolutionary information, implying that this may be within the realm of language models.^12–14^ To investigate this, we focused on ESM-1b, a Transformer masked language model trained over 250 million protein sequences.^5^ We computed the mean attention map, taking the mean of all attention maps generated by the model across all of its attention heads and layers (**Fig. 1**). We show that residues with high attention to the active site residues are significantly enriched for allosteric interactions. This is done in a zero-shot fashion with only a single amino acid sequence as input, indicating that these are relationships learned during the original masked language modeling task.

**Fig. 1:**
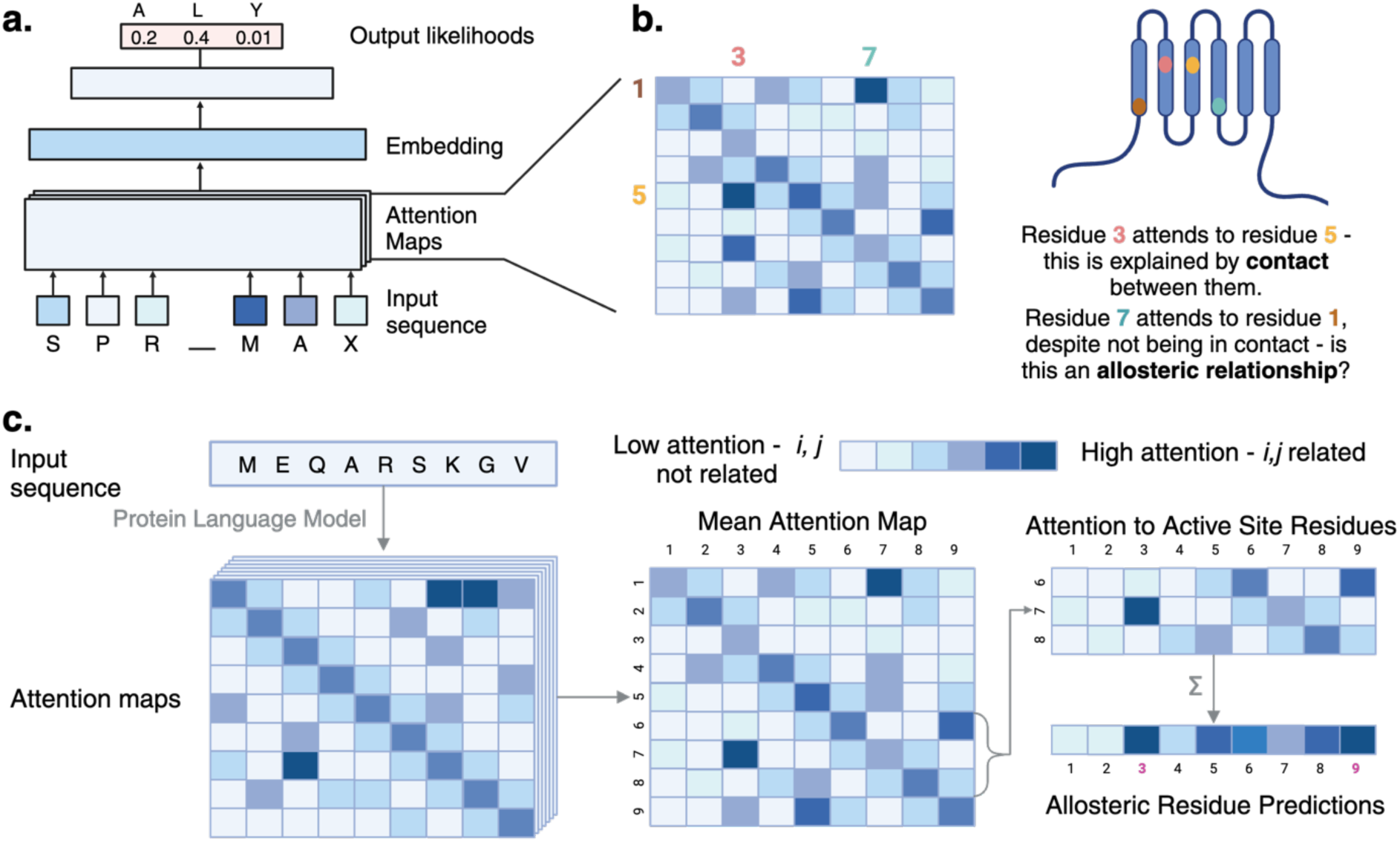
Spatially distant residue pairs highly attended to by language models may be allosterically related. (A) During masked language modeling, protein language models learn attention maps as representations of relationships between distant residues. (B) It is known that these attention maps sparsely learn structural contacts – if two residues are in contact, they may have high attention to each other. However, if they have high attention and are not in contact, we hypothesize that they may be related by allostery. (C) A protein sequence is input to the protein language model. Attention maps are extracted and the mean across all heads and layers is taken forward. The attention of all other residues to the active site residues is summed, and these are used as the output scores **(Methods)**.

We find that self-attention maps produced by language models from a single amino acid sequence identify residues in known allosteric sites substantially better than current methods. As a state-of-the art comparator, we study Ohm, a network model that takes a crystal structure as input. Ohm is used broadly in the literature to identify allosteric residues and communication pathways.^15^ As a sequence-only comparator, we study EVcouplings, a pipeline for sequence covariation analysis from multiple sequence alignments.^15,16^ Further, the language model correlates best with experimental deep mutational scanning and alanine scanning data mapping allostery in a GTPase and a GPCR, respectively. Strikingly, the language model can even discover meaningful allosteric relationships in structurally unresolved regions of proteins. More broadly, our study suggests that allostery is another emergent property of protein language modeling, which enables single-sequence allosteric residue discovery.

## Results

The architecture of a Transformer-based protein language model learns multiple “heads” of attention in each layer of the model.^8^ This is thought to allow each head to learn different information about the sequence.^8^ ESM1b and ESM2, the models studied in this work, have 20 heads per layer, and at least 33 layers. To derive a single attention map with information from all of these, we take the mean of the attention maps across the heads and layers. From this, we sum the attention of each residue to the active site residues, using this sum as the final per-residue score (**Fig. 1**). Any residues sequence-adjacent to an active site residue were excluded. We use the remaining values to rank residue-residue pairs for potential allostery (**Methods**).

### Attention maps from protein language models accurately identify allosteric sites

We first asked whether language models could identify residues binding allosteric modulators validated by crystal structures. We collated a dataset of 24 proteins – 19 from a previous 20-protein dataset used for benchmarking the network model Ohm, and five newly added.^15^ The set of active site or allosteric site residues were labeled from the crystal structures as those in contact with the substrate or allosteric modulator, respectively (**Fig. 2a**). The previous dataset was modified as described in ***Methods*** – removing a protein with subunits reported as separate UniProt IDs (ATCase), and including five new proteins to expand functional classes and types of binding partners (iGluR2/AMPA GluR2, MEK1, AR, AKT, IDH).

**Fig. 2:**
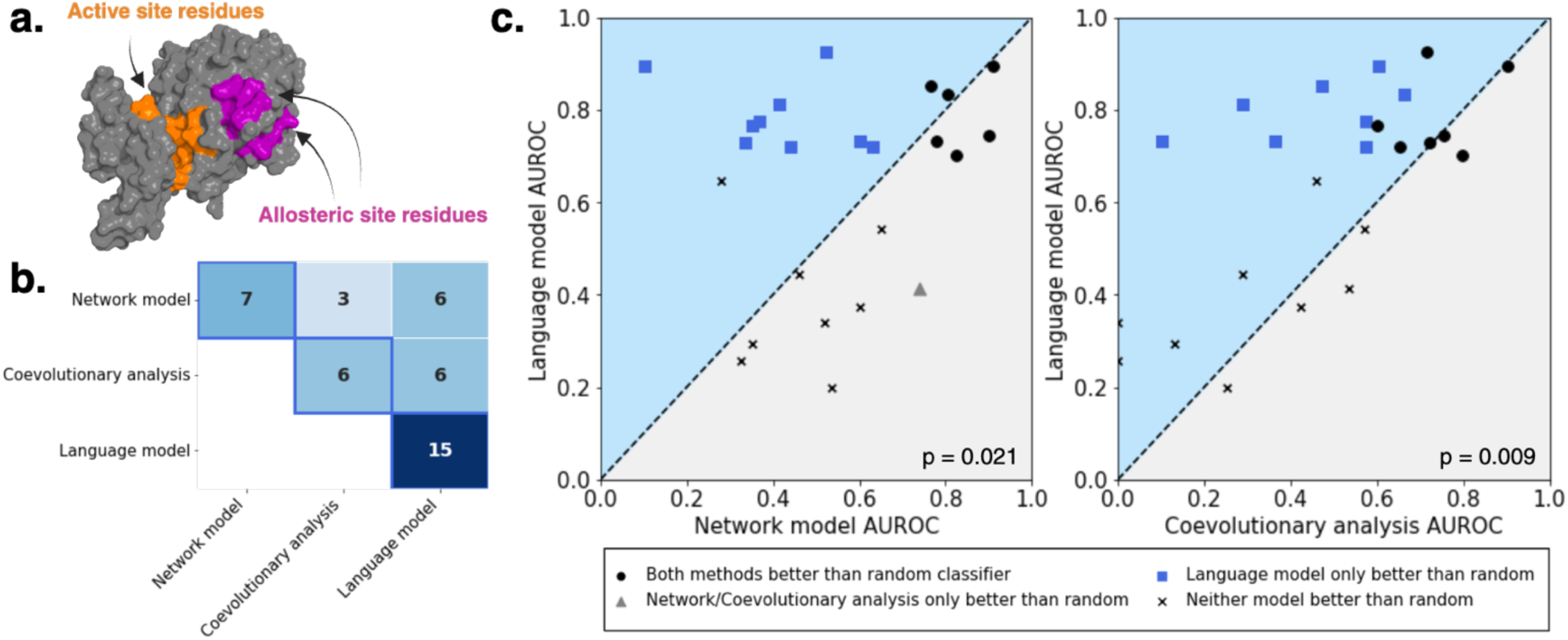
High-attention residue pairs identify allosteric relationships across diverse proteins. (A) “Active site residues” are defined as residues in contact with substrates, and “allosteric site residues” are defined as residues in contact with an allosteric modulator **(Methods)**. Modified PTP1b structure (from PDB:1PTY) used as example. (B) Correlation matrix of model predictions. Each cell contains the number of proteins with allosteric modulator contact residues classified better-than-random (“successes”) by both models, out of 24 total proteins in the benchmark set. The diagonal axis (outlined) contains the total number of successes of each model; for example, ESM-1b, a protein language model, had 15 successes. The language model successfully predicts allosteric residues for all but one of the Ohm (network model)-identified proteins and all of the EVcouplings (coevolutionary analysis)-identified proteins, in addition to 6 additional proteins that other models could not predict. (C) Language model ESM-1b accurately identifies allosteric residues in significantly more proteins than both other methods. AUROC is calculated per-protein for 24 proteins and plotted for each method, attempting to classify all allosteric residues in each protein. p-values are calculated with Fisher’s exact test comparing the number of proteins predicted better-than-random for each model by permutation testing. Proteins predicted better-than-random by both models are shown in bolded black circles, by ESM-1b shown in blue squares, by Ohm or EVcouplings shown in gray triangles, and by neither as black crosses.

The language model ESM1b performed well on this dataset, significantly outperforming the network model Ohm^15^ and the coevolutionary analysis pipeline EVcouplings^16^ with their default settings and maximum available MSA depths. ESM1b predicts allosteric sites in significantly more proteins across a benchmark set of 24 proteins (**Fig. 2b,c**). While Ohm was a statistically better-than-random predictor for 7/24 proteins, and EVcouplings for 5/24, ESM1b predicted the allosteric sites in 15/24 proteins better than random (p < by permutation testing) (**Fig. 2b, Table S1)**. On a per-protein basis, ESM1b generally scores highest as measured by AUROC (**Fig. 2c**). Eight of the proteins did not have sufficient MSA coverage at one or more active site or allosteric site residues for EVcouplings to fill a coupling matrix at those positions, rendering it impossible to make complete predictions for those proteins **(Fig. S1)**. However, enabled by the single-sequence-only requirement, the language model can make predictions at these residues, yielding higher accuracy across multiple such proteins (Fig. S1).

We also explored various ablations **(Fig. S2)** and modifications. Prediction of allosteric residues on this dataset did not scale linearly with model size; no significant improvements were seen from scaling beyond 150M parameters **(Fig. S3)**. We further explored single layers as predictors **(Fig. S4)**. It has been shown previously that early layers generally learn more surface-level, simple correlations, while deeper layers learn complex relationships.^17^ In general agreement with this, it appeared that deeper layers were somewhat better predictors than early layers (with layer 31 identifying the most allosteric sites), although accuracy on particular proteins varied widely between layers. Plotting the distance from the nearest active site residue against the attention score **(Fig. S5)**, we find that the language models identify high-attention residues across a range of distances from the active site in many cases. However, overall, high-attention residues tend to be enriched nearby in sequence to the active site residues.

It is likely that not all residues in contact with an allosteric modulator potentiate allostery.^18,19^ Therefore, instead of requiring that *all* allosteric modulator contact residues in the benchmark set are found, we evaluated when any given *one* residue is required to be found **(Fig. S7)**. This was represented by the percentage of total residues the model would have to search through to have correctly found a single residue in contact with the known allosteric modulator – i.e., a depth of 0.1 means the model scores 10% of the protein higher than the highest-scoring residue in the known allosteric site. When evaluated in this lens, the language models show the highest density of low-depth proteins of the three models, performing slightly better than Ohm and significantly better than EVcouplings **(Fig. S7)**.

### Attention maps identify allosteric relationships in diverse proteins

Having established that language models can accurately identify allosteric residues in the aggregate, we next sought to explore several diverse examples from the benchmark set to investigate performance on important enzyme classes. We selected five human proteins to investigate further:

- Kinase (PDK1, 3-phosphoinositide-dependent protein-kinase 1)
- Dehydrogenase (IDH1, isocitrate dehydrogenase 1)
- Hydrolase (CASP1, caspase 1)
- Transferase (ATPS, ATP Sulfurylase)
- Phosphatase (FBP, fructose bisphosphatase)

On the first three, the language model outperformed both the network model and the coevolutionary analysis (**Fig. 3, Fig. S11)**, accurately identifying the allosteric site at the dimer interface of IDH1 and allosteric residues in known the CASP1 network. However, on ATPS, none of the tested models performed well, while on FBP, Ohm performed the best. These five proteins represent a range of symmetries, sizes, functions, and allosteric site arrangements – in cases such as PDK1, the allosteric site is nearby the active site compared to caspase 1 or ATPS. We briefly explored these five enzymes, attempting to understand model behavior across a range of enzyme classes. In-depth descriptions of these enzymes, active site labels, and their data are available in the supplementary text.

**Fig. 3:**
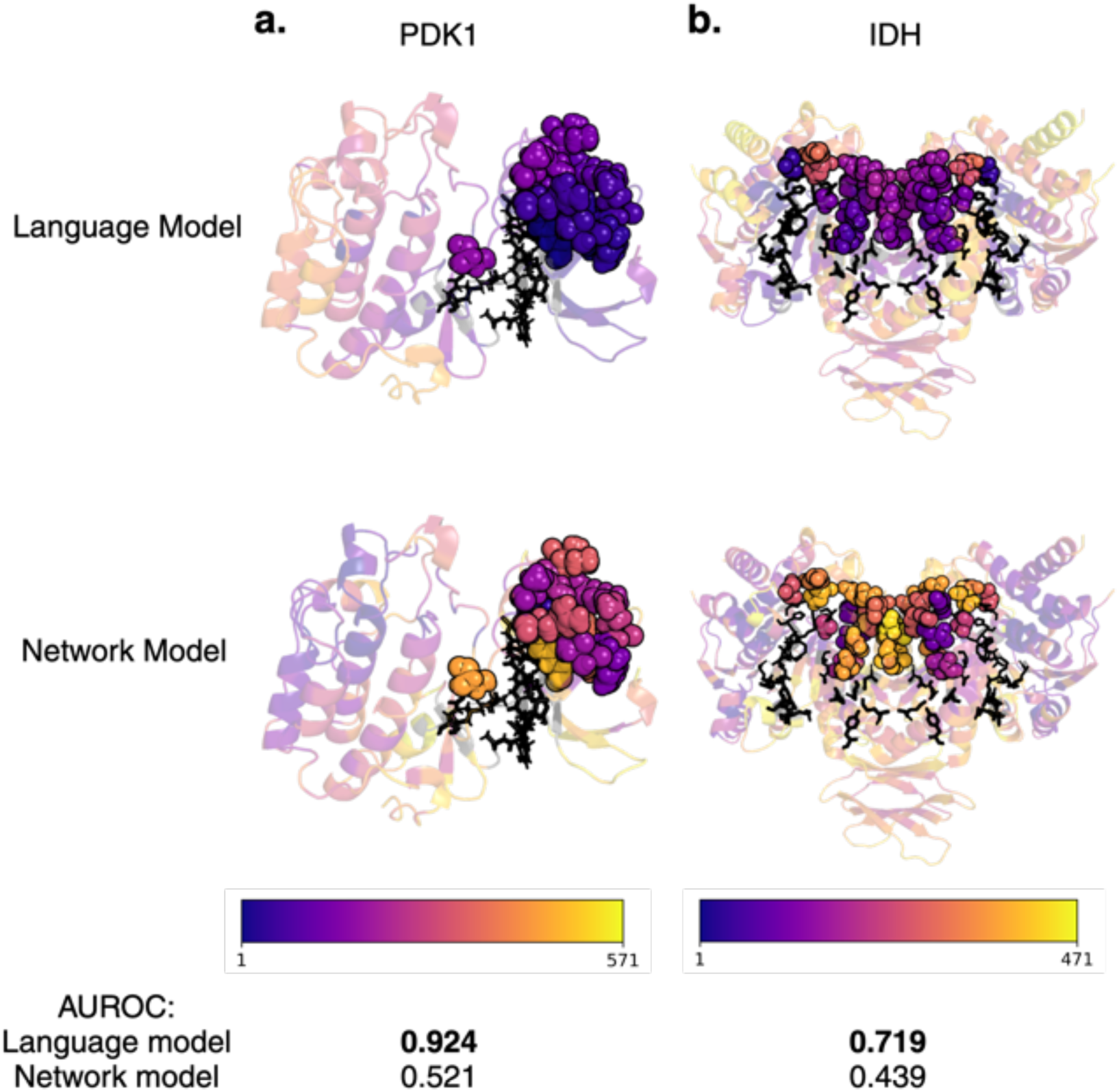
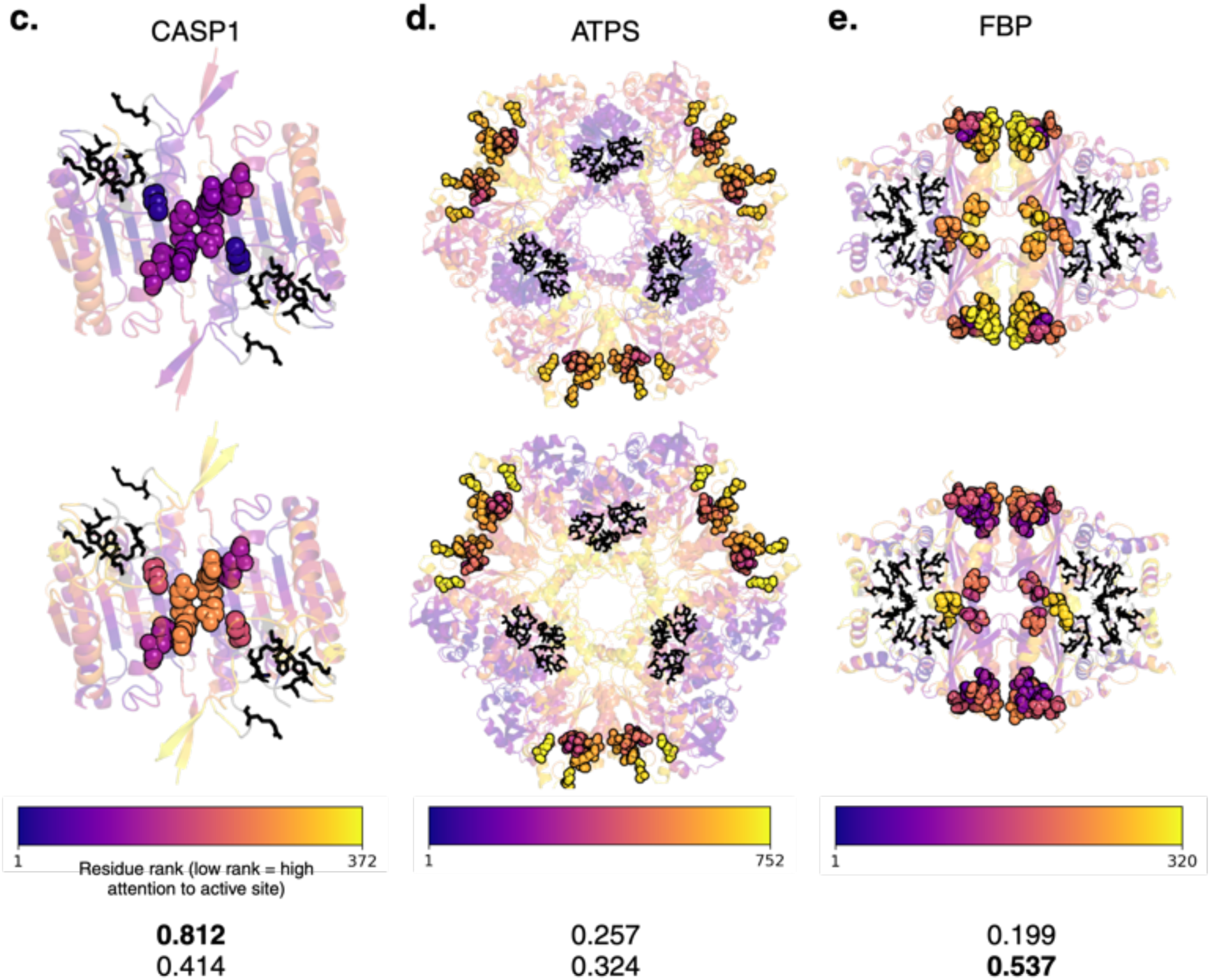
Protein language model attention accurately identifies known allosteric sites across a variety of protein classes. Five proteins are selected from the 24-protein benchmark set as examples – these include three for which the language model performs best, one for which both the network model and the language model perform poorly, and one for which the network model performs best. Bar plot for all proteins is shown in **Fig. S11.** (A-E) Structures of example proteins. Crystallographically resolved residues are colored on ribbon diagrams by rank-scored attention or Ohm score, with dark purple being highest-attention (or Ohm score) to active site residues. Allosteric residues (in contact with an allosteric modulator) are represented and colored as spheres. Active site residues (in contact with a substrate) are represented in sticks, outlined in black and colored in orange. ESM1b-colored structures of all proteins are shown in **Fig. S12.**

As language models only take in a single sequence as input, with no structural requirements, they may enable prediction of structurally unresolved allosteric relationships. This is a capability that has, to our knowledge, not been demonstrated by alternative allosteric residue prediction methods. We investigate this by probing allosteric relationships between the DNA-binding domain (DBD) and ligand-binding domain (LBD) in the androgen receptor, a therapeutically relevant nuclear receptor. This protein is incompletely resolved with X-ray crystallography and has comparatively little sequence homology for MSA generation **(Fig. S7a)**. It is known that anti-androgen binding in the LBD can reduce DNA-binding directly, despite these domains being sequence-distant.^20–23^ However, the mechanism underlying this interdomain relationship is unknown. We find four residues in the DBD with high attention (>90^th^ percentile overall) to the androgen-binding site in the LBD, including a zinc-binding residue **(Fig. S7b)**. Three of these residues are mutated in patients with androgen insensitivity syndromes. Two of these three variants are known biochemically to ablate androgen-binding.^24–27^ The identification of these residues as potentially allosteric could provide an explanation for an interdomain relationship driven by changes in zinc-binding and DNA-binding specificity, supporting the notion in literature that the LBD and DBD may be allosterically linked.^20,23,28^ A more in-depth explanation of this investigation is available in the supplemental text.

### Attention maps identify binding allostery over unfolding effects in K-Ras

Effects of active-site-distant mutations on protein function can be mediated both by allosteric coupling between the active site and the mutated residue, or by mutation-induced misfolding/unfolding. We wondered whether the language model was simply identifying residues that would generally unfold the protein, or whether meaningful allosteric relationships were reliably being identified. Recent deep mutational scanning methods, validated by targeted mutagenesis and calorimetry, have been developed to deconvolute allosteric effects of mutations on free energies of binding from general effects on free energies of folding.^19,29^ These represent datasets labeled experimentally at every residue, compared to the positive-unlabeled datasets from the previous crystallography benchmark set **(Methods)**. This method has been applied to full-length K-Ras, and an additional three short protein-binding domains – in this analysis, we focus primarily on K-Ras as it is the only available full-length protein (188 residues), with a brief analysis on the comparatively much shorter binding domains (55-84 residues).^19,29^

Deep mutational scanning was used to probe allostery in K-Ras, a therapeutically relevant oncoprotein, deconvoluting the contributions of allosteric residues throughout the protein on binding to each of six binding partners (RAF1, PIK3CG, RALGDS, SOS1, DARPin K27, DARPin K55). Using the definitions for binding residues and allosteric residues from ref. 19 **(Methods)**,^19^ we ask whether, across multiple binding partners, the language model (1) accurately identifies deep mutational scanning-validated allosteric residues, (2) correlates differently with binding effects and folding effects of mutations, and (3) identifies the known allosteric sites (physiological GTP+Mg^2+^ binding site, and therapeutic sotorasib binding site). For each case, we look at the attention of each non-binding residue in K-Ras to the residues contacting each respective binding partner.

We find that language model attention maps identify allosteric residues in K-Ras remarkably well; high-attention residues are strongly enriched for allosteric residues (**Fig. 4a**). There were >6 allosteric residues among the top-10 highest attention residues for five of the six binding partners (hypergeometric p-value ≤ 0.005 for all six). In comparison, the top-10 scoring residues by the coevolutionary analysis (EVcouplings) were significantly enriched for one of the six binding partners, and by the network model (Ohm) for zero of the six. The average AUROC across all binding partners was 0.86, compared to 0.63 for the network model and 0.68 for coevolutionary analysis **(Fig. S8a,b)**.

**Fig. 4:**
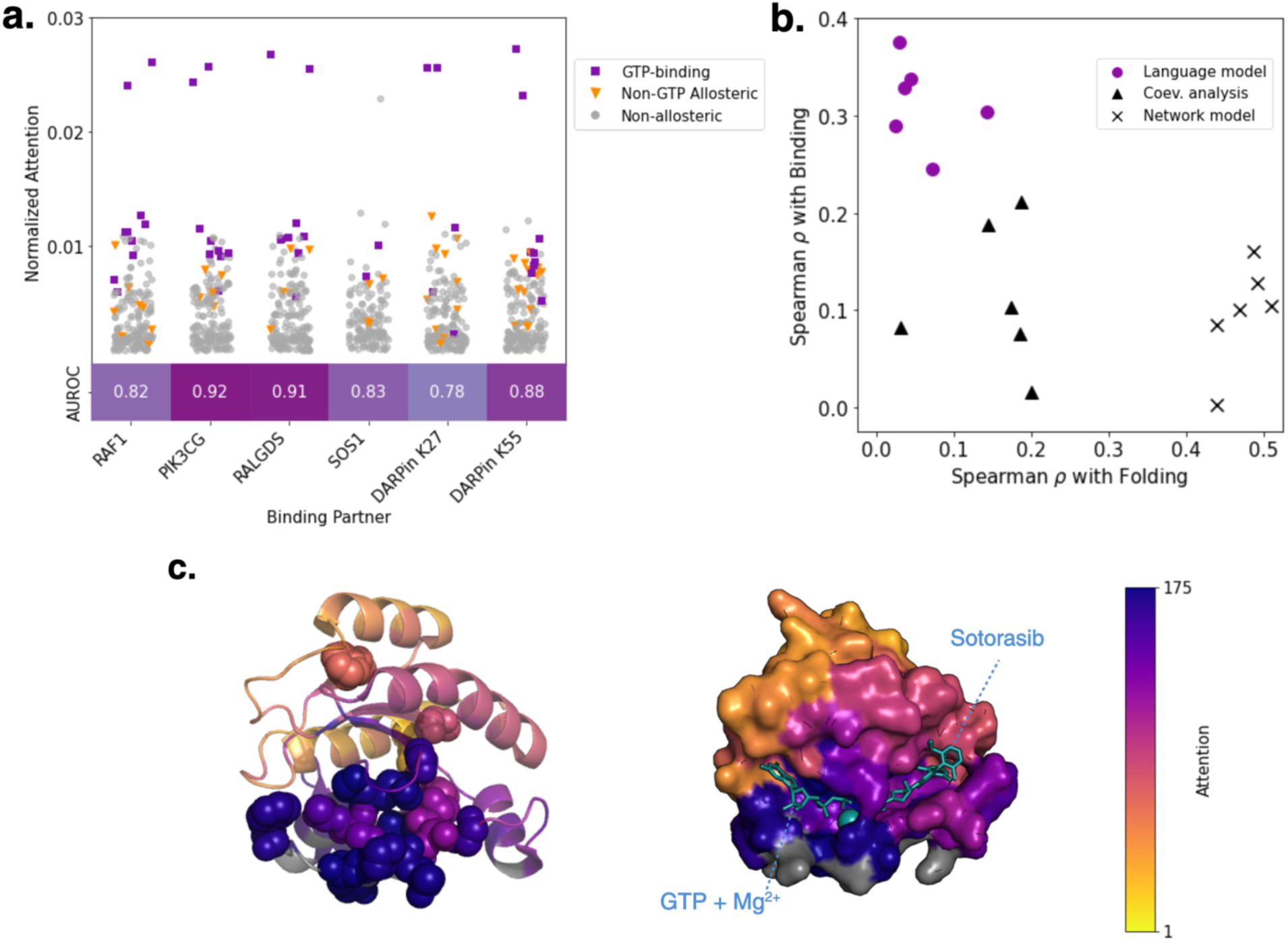
Protein language models accurately identify allosteric residues from a deep mutational scan of K-Ras, correlating with allosteric binding effects but not unfolding effects. Deep mutational scanning methods have been developed to differentiate allosteric effects of mutations on protein-protein binding from general effects on folding.^19,29^ We search for residues with high attention in self-attention maps to the binding residues for each binding partner (RAF1, PIK3CG, RALGDS, SOS1, DARPin K27, and DARPin K55). Binding residues and allosteric residues are as defined in ref. 19, as are binding/folding free energy changes; including *all* mutagenesis-identified allosteric residues, not only those in modulator binding sites.^19^ (A) Top: Scatter plot of attention to the binding site of each binding partner (normalized by number of binding site residues). Allosteric residues in the GTP-binding site are colored purple, outside the GTP-binding site (including the sotorasib-binding site) orange, and non-allosteric residues gray. Allosteric residues are highly enriched among the top-10 highest attention residues, with hypergeometric p-value <0.005 for SOS1 and <10^-4^ for all others. Bottom: AUROCs predicting allosteric residues for each binding partner. Average AUROC across all binding partners is 0.86, and all are significant with p < 0.005 by permutation testing. (Comparisons with network model and coevolutionary analysis in **Fig. S8a,b**). (B) Spearman correlations of each model with absolute mean ΔΔG_folding_ and absolute mean ΔΔG_binding_ as calculated from data in ref. 19.^19^ Purple dots represent language model attention scores, triangles represent coevolutionary analysis scores, and crosses represent network model scores. Language model significantly correlates with binding (p < 0.005), but not folding, for all 6 binding partners. Coevolutionary analysis correlates with binding for 2/6 proteins (p<0.05), folding for 2/6 proteins (p < 0.05), and neither for the remaining 2 proteins. The network model correlates with folding very strongly for all 6 binding partners (p < 0.001), and does not correlate significantly with binding. (Full data in **Fig. S8c**) (C) (PDB: 6OIM) Structure of K-Ras with residues colored by rank-scored attention to RAF1 binding site. Left: All experimentally determined allosteric residues shown as spheres.^19^ Right: Matched pose, with all residues colored according to attention, allosteric modulators GTP, Mg^2+^, and sotorasib shown with sticks colored in cyan, and binding residues colored in gray.

When we calculate Spearman correlations of attention to binding-site residues with the free energy changes (ΔΔG) of binding and folding as experimentally determined in Ref. 19, across all six binding partners (and largely distinct definitions of binding sites), the correlations are significant (p < 0.005) with binding, but not with folding (**Fig. 4b**). In comparison, the network model and evolutionary couplings both bias comparatively more towards folding; the network model correlated strongly with folding ΔΔG for all binding partners, but not with binding ΔΔG, while coevolutionary analysis correlated with binding for two of the six partners, and folding for two of the six partners **(Fig. S8c)**.

Allosteric residues in both the physiological GTP/Mg^2+^-binding site and the small-molecule sotorasib binding site are identified accurately by the language model (**Fig. 4c**). The GTP-binding site, in particular, is identified across all six binding partners among the highest-scoring residues. Additionally, in the separate DMS dataset of three short protein binding domains, we find that self-attention correlates well for two of the three with the number of allosteric mutations at a particular residue **(Fig. S9)**.^29^ In total, within the context of a comprehensive deep mutational scan of allostery, self-attention is able to accurately identify allosteric residues and uniquely selects for binding allostery while largely avoiding residues that primarily induce unfolding.

### Attention maps identify allosteric sites and switches in a model GPCR

G-protein coupled receptors are rich in allosteric regulatory mechanisms and represent one of the most therapeutically exciting classes of proteins, targeted by ∼35% of drugs.^30^ Recent mutational scanning efforts aim to deconvolute allosteric signaling within the prototypical GPCR, beta-2 adrenergic receptor (B2AR). Heydenreich et. al. mutate nearly all residues in an alanine scan to identify the effects of each residue on signaling potency (measured as EC_50_) and efficacy (as measured by the amplitude of the receptor conformational change) when stimulated with the activator adrenaline.^18^ They identify a set of pharmacologically important residues that, when mutated, shift receptor signaling from wild-type-like to low-potency, low-efficacy, or both.

We asked whether attention to the adrenaline-binding site could be used to differentiate between allosteric and non-allosteric residues. We find that this is indeed the case; mutating residues with high attention to the adrenaline-binding site often diminishes potency, efficacy, or both (**Fig. 5a**). Of the top 20 highest-attention residues, 13 are allosteric as determined in ref. 18 (11 pharmacologically important, and 2 of 3 total low-expressing mutated residues that are part of known allosteric motifs) – a hit-rate of 65%, and a ∼4.5-fold over-enrichment of allosteric residues compared to random guessing (hypergeometric test p < 1x10^-7^).^18^ High attention correlates significantly with changes upon mutation in both logEC_50_ and amplitude, whereas Ohm and EVcouplings scores do not **(Fig. S10a,d,g)**. Among the top 50 highest-attention residues, there is a ∼2.6-fold over-enrichment compared to random guessing (31/50 for the language model vs. ∼12/50 expected by random guessing, hypergeometric p < 1x10^-9^), compared to a ∼1.4-fold enrichment for Ohm (17/50, hypergeometric p = 0.06) and a ∼1.6-fold enrichment for EVcouplings (19/50, hypergeometric p = 0.01) **(Fig. S10b,e,h)**.

**Fig. 5:**
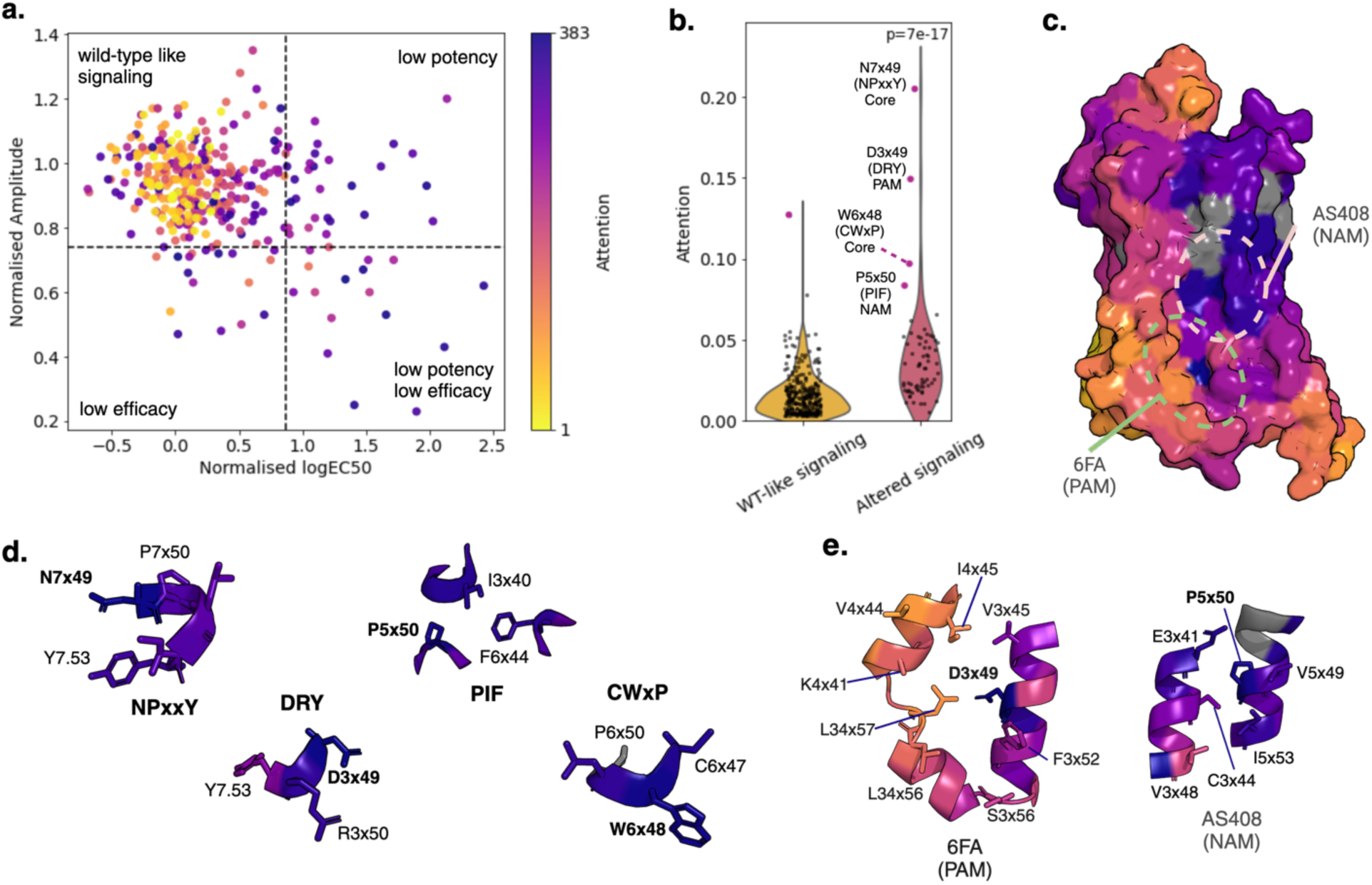
Protein language models identify allosteric residues in GPCRs, including microswitches and modulator-binding sites. (A) Residues in B2AR colored by rank-scored attention, as calculated by the language model ESM1b, to the adrenaline-binding site residues (as defined in ref. 18). Darker coloring indicates higher attention to the active site. Residues are plotted by potency (logEC50) and efficacy (amplitude) of signaling when mutated to alanine in ref. 18.^18^ Mutant residues that drive non-WT-like signaling (potency or efficacy) are treated as allosteric. Thresholds as in ref. 18 are marked by dotted lines, and non-WT-like mutants are treated as allosteric. Allosteric mutations are significantly enriched among high-attention residues (see also, **Fig. S10b**). (B) Residues grouped by pharmacological importance (ref. 18) and plotted by language model attention to the adrenaline-binding site. Positions which drive non-WT-like signaling when mutated have significantly higher attention to adrenaline-binding residues than WT-like positions (two-sided t-test p < 10^-17^). The top 5 highest-attention residues are colored in pink. Four of these (labeled) are part of well-described allosteric regulatory motifs, with one in each of the four known state-stabilizing “microswitches” (as labeled in refs. 18, 30).^18,31^ (C) (PDB: 4LDO) Structure of B2AR with surface-exposed residues by attention to active site, rank-scored by rank in structurally resolved residues only. Residues sequence-adjacent to active site are shown in gray. Color scheme is identical to Fig. 5A; darker residues have higher attention to active site. AS408 (NAM) binding site and 6FA (PAM) binding sites are circled in peach and green. Language model identifies two surface-exposed allosteric sites on B2AR that, when mutated, impact signaling. Each of these sites has at least 1 residue in the top-20 highest attention residues overall. Complete views of structures, and comparisons to Ohm and EVcouplings, are shown in **Fig. S10**. (D) Full structures of all four Class A GPCR allosteric microswitches (as labeled in refs. 18,30) with residues colored by attention to adrenaline binding residues as in Fig. 5A.^18,31^ Conserved residues labeled with numbering. All four microswitches are in the top-5 highest-attention residues. (E) Structures of allosteric modulator binding sites, with contact residues shown in stick representation and colored as in Fig. 5A.

Class A GPCRs such as B2AR are known to contain motifs that are involved in signaling and stabilize the active or inactive states. In particular, four such “microswitches” (NPxxY, DRY, PIF, and CWxP) that change orientation between active/inactive states and drive altered signaling when mutated are well-described in literature among Class A GPCRs.^18,31–34^ Among the top-5 highest-attention residues to the adrenaline binding site, the language model identifies four residues in these microswitches; one in each site (**Fig. 5b,d**). Among the top-20 highest-attention residues, the language model identifies 8 microswitch residues (out of 12 total microswitch residues) **(Table S2).** In comparison, Ohm identifies no microswitch residues among the top 20 scoring residues, and EVcouplings identifies two residues in the microswitches among the top 20 scoring residues **(Fig. S10m,n,o)**.

B2AR has small-molecule binding sites for a negative allosteric modulator (NAM) and a positive allosteric modulator (PAM). We asked whether residues in these known binding sites were identified by the language model as allosterically linked to the adrenaline-binding site. Indeed, in both sites, at least one residue is in the top-5 highest attention residues to the active site (**Fig. 5d,e, Table S2)**. D3x49 of the DRY motif and P5x50 of the PIF motif are present in the PAM and NAM binding sites, respectively. Ohm identifies residues in the PAM-binding site with a high coupling intensity to the adrenaline-binding site, while EVcouplings identifies residues in the NAM binding site. Only the language model identifies residues in both sites as high-scoring **(Fig. S10c,f,i)**.

## Discussion

We show that attention maps from protein language models can, in a zero-shot fashion with a single wild-type sequence and no input structural information, identify allosteric relationships within a protein at single-residue resolution. This performs better than state-of-the-art methods on a benchmark set of allosteric sites, a deep mutational scanning dataset, and an alanine-scanning mutational dataset. We observe these effects across diverse proteins by class, fold, function, size, and symmetry, and probe potential allosteric communication in a structurally unresolved protein. We show that, in the case of a comprehensive deep mutational scan of allostery in K-Ras, the language model accurately identifies allosteric residues and correlates well with mutational effects on binding allostery, while avoiding general effects on unfolding. Further, we show that allosteric regulation in a prototypical Class A GPCR, the beta-2 adrenergic receptor, can be identified accurately by the language model, including identification of critical residues in known allosteric modulator binding sites and microswitches. The sequence-only nature of language models enables these predictions to be made in the absence of structure or a multiple sequence alignment, potentially expanding the analysis of allosteric communication to the >98% of proteins without reported high-resolution structures.

Despite the useful information learned by language models, allosteric residue discovery remains difficult. That protein language models can learn information about allostery from evolutionary information and, in doing so, outperform current tools, sets the stage for further language model-based methodological improvements. Small scale supervision or finetuning could help improve prediction performance further. It is also likely that allosteric connections through an intermediary are being missed – if *i* affects *j*, which affects active site residue *k*, this may not be perceived when just looking at outgoing attention from *i*. We also find it interesting that a single sequence was sufficient to predict allosteric relationships in some proteins that form multimers, indicating that the model may not need explicit information about an interface to discover allosteric couplings, though better methods of representing symmetry as input to the model could lead to performance improvements on multimers. Structure-informed language models have shown the ability to navigate complex sequence-structure-function landscapes^4^, and could yield significant general improvements on sequence-only models for allostery prediction.

## Methods

### Identifying allosteric residues from attention maps

Given a model with *N* layers, *K* heads per layer, *i* = 1, …, *N* layer index and *j* = 1, …, *K* head index: let 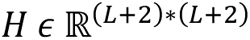 be an attention map produced by the model at every head. We define *H*^(*i,j*)^ to be an attention map associated with head *j* at layer *i*. We take a mean over all heads in each layer, creating a global mean attention map 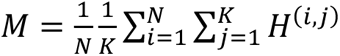.

Now let *l* = 1, …, *L* + 2 be the index of a row in *M*. Take 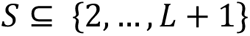 to be the set of active site residues associated with row *l*. We then create a submatrix *M_s_* to be a matrix consisting of only the rows of *M* indicated in *S*.

For example, for set 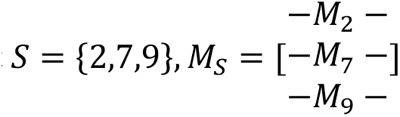. To get per-residue allosteric coupling score 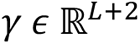, sum along rows of 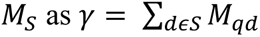 where *M_qd_* is a specific row of *M*.

Residues sequence-adjacent to active site residues were removed. Remaining residues were then ranked by score, with the highest-scoring residue taken as most likely to be allosteric as predicted by the model. The ranks and scores are then used for scoring on the benchmark set, deep mutational scanning, and exploratory analyses.

### Benchmark set

24 proteins with known allosteric modulators and substrates were chosen as a benchmark set. 19 proteins were used from Wang et. al., supplemented with additional manually curated information about active sites and allosteric sites, and 5 additional proteins were added for diversity across protein functional classes.^15^ Proteins were manually selected from the ASbench website and manually supplemented with complete active site/allosteric site information, with requirements that (1) a substrate-containing PDB file and allosteric modulator-containing PDB file, both with the same UniProt ID, were available for all identified substrates and modulators binding at unique sites, (2) protein length <1024 residues, (3) full-length sequence reported as a single UniProt ID, and (4) reported allosteric site does not overlap with active site.^35^ Aspartate transcarbamoylase was removed from the Wang et. al. dataset as the sequence is split across two separate proteins with unique UniProt IDs. The added proteins include: AKT, a master regulator serine/threonine kinase with a promiscuous binding site and peptide substrate, Androgen receptor, a multi-functional signaling molecule and transcription factor, iGluR2/AMPA GluR2, a glutamate-sensitive ion channel, MEK1, a dual-specificity kinase that is allosterically inhibited in its BRAF-bound state, and IDH1, an oxidoreductase with plasticity in the allosteric site allowing a wide range of allosteric modulators.^36–38^ All added proteins are therapeutically relevant, with approved or clinical-stage drugs.

All known active sites with crystal structures of bound substrates were included for each protein, along with the set of known allosteric sites with crystal structures of bound modulators. PDB files were renumbered according to the corresponding Uniprot sequence. Residues with an atom within 3.4A of a modulator or substrate atom were considered in contact with the modulator or substrate. Supplemental Table S3 lists active site and allosteric site structures and modulators used for this approach. Multiple ligands may exist per site – the union of contact residues was taken as that site. Residues within 1 sequence position of an active site residue were explicitly not allowed to be labeled allosteric. These 24 proteins are similar to a positive unlabeled dataset; that is, we can say that the residues labeled as positive are most likely positive, but all other residues could potentially be positive OR negative. Therefore, AUPR and AUROC are likely higher than predicted here, as unlabeled data is treated as negative for statistical analysis here. Within the context of each protein, AUROC was used to score models as classifiers of allosteric residues, where p < 0.05 by permutation testing was taken as a better-than-random prediction for each protein (a “success”). Fisher’s exact test was used to compare success/failure rates across models.

### Network model

All 24 proteins were run on the Ohm web server with default settings. The active site PDB structure with Uniprot-matched numbering was taken as input. The active site residues (within 3.4Å of substrates) were individually selected in each case, and the allosteric site ligand was removed for structures containing it in the active site PDB. For proteins with multiple active site structures, Ohm was run on all structures independently and the highest-performing structure as measured by AUROC was taken as the network model score for that protein. The ‘ACI’ file (allosteric coupling intensity) was downloaded and processed identically to the attention values. Unmodeled/missing allosteric residues in the structure were treated as impossible to find/not found.

### Coevolutionary analysis

Uniprot codes for all proteins were input to the EVcouplings server. The best MSA for each protein as marked by the server was used to move forward. The coupling matrices between all residues were processed identically to the attention values, with sequence-adjacent residues set to 0. For 8 proteins, the MSA did not sufficiently cover active site residues or allosteric site residues and EVcouplings was unable to make predictions at those positions. Predictions were made using the available active site residues.

### Mutational scanning

Potency and efficacy data as well as WT-like/non-WT-like signaling labeling from ref. 18 was used for B2AR analysis. Adrenaline contact residues were taken as “active site” residues. Definitions of microswitches/motifs and PAM/NAM sites are used as in refs. 18 and 30.^18,31^

K-Ras analysis was based on experimental data from ref. 19.^19^ Binding and folding ΔΔG were calculated as the difference between a single mutation’s ΔG and the WT ΔG value. The mean absolute ΔΔG at each position was taken forward for analysis. Labeling of binding site/allosteric site residues was used as in Ref. 19 for calculation of AUROCs. PDB:6VJJ was used as input for Ohm (default settings), and the sequence Uniprot:P01116-2 was used as input for the language model and EVcouplings (default settings).^39^ PSD95-PDZ3, GRB2-SH3, and GB1 analysis was based on experimental data from ref. 29, with binding site residues defined as in ref. 29.^29^ Spearman correlations between the scores output from each model and the number of allosteric mutations at each site, as labeled in ref. 29, were calculated. PDBs:1BE9, 2VWF, and 1FCC were used as input for Ohm for PSD95-PDZ3, GRB2-SH3, and GB1 respectively.^40–42^

## Acknowledgements

We would like to thank Onn Brandman and the members of the Kim lab for helpful discussions on this project.

## Author contributions

Conceptualization, methodology, interpretation: all authors; Computational experiments and software development: G.R.K.; Writing: all authors.

## Supplementary Text

### Attention maps identify allosteric relationships in diverse structurally resolved proteins

Kinases are among the most interesting allosteric proteins therapeutically. Synthesis of selective ATP-competitive inhibitors has historically been difficult due to the conserved nature of the ATP-binding site, leading allosteric drug design to be particularly interesting in the class.^43^ A number of high-profile allosteric kinase inhibitors have been approved in recent years for targets like MEK1/2 and BCR-ABL1 leading to renewed interest. To explore whether we could accurately identify allosteric residues in therapeutically interesting kinases, we explored PDK1 – one such disease-implicated kinase included in the crystallography benchmark set. PDK1 is a serine/threonine kinase that plays a crucial role in cell growth and proliferation. In this case, we include the ATP-binding site as the active site. There are two allosteric sites – one binding the fragment activator 2A2, and another binding a peptide activator (the PIF pocket). In this case, we identify residues contacting the fragment activator or the small-molecule PIFtide mimic RS1 as allosteric. We find that the protein language model is able to accurately identify both allosteric domains in PDK1 (**Fig. 3a**). Most of the residues in contact with RS1 show up as high-attention (dark blue/purple), indicating that a large part of the binding site may be involved in allosteric relationships with the black active site residues. The second allosteric site binding 2A2 is also identified as high-attention. The identification of both known allosteric sites – the PIF pocket binding RS1 and the 2A2-binding site – highlights the utility of protein language models to accurately identify allosteric sites on monomeric kinases for small-molecule drug development.

Isocitrate dehydrogenase (IDH) plays a crucial role in the Krebs cycle, a fundamental part of cellular respiration, by catalyzing the oxidative decarboxylation of isocitrate to alpha-ketoglutarate. Mutations in isocitrate dehydrogenase are linked to various types of cancer, highlighting its importance in normal cellular metabolism and the potential consequences of its dysfunction.^44^ We choose the NADPH binding site, the isocitrate binding site, and the magnesium cofactor binding site as the active sites. The allosteric sites include the citrate binding site and a small-molecule binding site, which is capable of changing conformation to take on a variety of small-molecule modulators.^38^ While PDK1 is a monomeric enzyme, IDH1 exists as a homodimer, and is allosterically regulated by binding both within each subunit (by citrate) and at/around the subunit interface (by small molecules) – allowing us to explore the language model’s ability to identify allosteric sites at these interfaces. No particular steps were taken to deal with the symmetric nature of IDH1; the model simply took in the sequence of a single subunit. We find that, even without the knowledge that symmetry or an interface exists, the language model identifies both allosteric sites better than the network model (**Fig. 3b**). The language model identifies Ser326 in the allosteric citrate binding site as a high-attention residue; S326P has been found in high-grade gliomas as reported in ClinVar, but has unknown significance.^45^

Caspase-1 is an allosterically regulated protease involved in apoptosis and inflammation. Two asymmetric monomers form a dimer, which then arranges symmetrically to form a tetramer (dimer of dimers). While targeting the active sites (sitting at the outside edges of the tetramer) with synthetic inhibitors remains difficult, success has been seen with allosteric inhibition (at the central interface of the tetramer). As the active site, we choose the gasdermin mimic peptide cleavage sites. The allosteric site in this case is the binding epitope for small-molecule inhibitors. Despite having no knowledge of structure or symmetry, the language model identifies the allosteric site at the interface more accurately than the network model (**Fig. 3c**). While the network model identifies some residues surrounding the allosteric modulator contacts, it fails to score allosteric modulator contact residues themselves highly. Additionally, the language model identifies the asymmetric dimer coordinating residue Cys391 as high-attention to the active site; this residue can be allosterically trapped, and mutating it abates the effect of inhibitors binding at the central interface.^46,47^

Having now explored a monomeric kinase, a dimeric oxidoreductase, and a tetrameric hydrolase, we turn our attention to ATP Sulfurylase (ATPS) – a large enzyme with six identical subunits stacked into a circular hexamer. ATPS catalyzes the first step in the sulfate activation pathway, a crucial reaction in the synthesis of sulfur-containing compounds in organisms. ATPS is allosterically regulated by PAPS, an AMP-derivative. ^48^ We define the active site as the ATP-binding pocket. Both the language model and the network model perform poorly in attempting to predict the PAPS binding site (**Fig. 3d**). The language model false positives skew nearby to the active site residues – in comparison, the network model false positives are at the far edges of the protein.

We find another example of this sequence-distance dependence of attention, albeit to a lesser degree, in the enzyme Fructose 1,6-bisphosphatase (FBP). FBP is a critical metabolic enzyme that catalyzes the conversion of fructose-1,6,bisphosphate to fructose-6-phosphate, requiring a metal ion as a cofactor. It is allosterically regulated by AMP and synthetic allosteric inhibitors.^49,50^ In this case, the active site is defined as the fructose-6-phosphate binding site and the magnesium cofactor binding epitope. The AMP-binding site is notable as it is particularly distant from the substrate binding site. When FBP is analyzed by the language model, a few of the residues in the known allosteric site are highlighted as high-attention residues compared to the surrounding residues – but most residues in the site have low attention to the active site. In comparison, the network model scores nearly all of the surrounding residues more highly, including the secondary allosteric site at the dimer interface (**Fig. 3e**) – indicating that, for this case, the network model clearly outperformed the language model.

### Attention maps probe allosteric relationships in a structurally unresolved protein

To this point, we have explored proteins and protein domains that are structurally resolved. Notably, however, protein language models potentially enable prediction of structurally unresolved allosteric communication. This is a capability that has, to our knowledge, never been demonstrated by alternative allostery prediction methods. We investigate this by attempting to answer an open question about the androgen receptor (AR), one of the proteins in our benchmark set. The ligand-binding domain (LBD) and DNA-binding domain (DBD) of AR were thought to function largely independently; androgen binds the LBD, inducing a local conformational change that exposes the nuclear localization sequence.^51^ The DBD dimerizes, translocates to the nucleus, and binds DNA, activating transcription.^52^ Experimental evidence has shown, however, that anti-androgen binding in the hormone binding site can ablate DNA-binding directly – despite a large sequence-distance between the ligand-binding domain (LBD) and the DNA-binding domain (DBD).^20,23,53^ Additionally, clinically observed mutations in the DBD have been reported to ablate androgen binding.^25,26^ Recent cryo-EM work has shown that there is interdomain allostery mediated by contact surfaces between the LBD and DBD in the DNA-bound state.^23,28^ The extent and mechanism of this interdomain allostery and communication is unknown.

Androgen receptor has no full-length crystal structures reported in the PDB, and no crystal structures including both the DBD and the LBD **(Fig. S7a)**, making it difficult to use a structural network model like Ohm to make cross-domain predictions. Moreover, there is comparatively little sequence homology to generate an MSA from, making it difficult to use a deep MSA for sequence covariation with EVcouplings. We wondered whether treating the androgen-binding site as the active site would allow us to identify allosterically linked residues in the DBD.

The language model identifies four locally high-attention residues (>90^th^ percentile) in the DNA-binding domain of interest **(Fig. S7b)**; C615, K619, C620, and F584. Three of these are clinically observed disease-associated variants. C615, K619, and C620 are not present in any reported crystal structure; the analogous residue for F584 is present in a single rat androgen receptor DBD structure (PDB:1R4I). The DBD is separated from the LBD by a hinge region that is not structurally resolved crystallographically or by AlphaFold **(Fig. S7a)**. Of these, the mutation C620S is known biochemically to ablate ligand binding and transactivation, and leads to partial androgen insensitivity syndrome.^26,27^ F584 deletion is a clinically observed mutation that results in complete androgen insensitivity syndrome; while it is directly responsible for DNA-binding, F584 deletion also ablates androgen binding, an effect which remains unexplained to our knowledge.^25^ C615Y, a zinc-coordinating cysteine mediating DBD dimerization, was found in a patient with complete androgen insensitivity syndrome.^26^ K619 remains undescribed from the perspective of allostery or testosterone binding, but its acetylation is known to be required for hormone-dependent nuclear localization.^24^ The identification of these residues provides a potential explanation for an anti-androgen/DNA binding relationship. Anti-androgen binding may limit the ability of C615 to bind zinc and/or F584 to mediate ARE-binding specificity, affecting dimerization and/or DNA-binding. This supports the notion that the LBD and DBD may be allosterically linked, and provides a potential explanation for the potency of modulators and single-amino-acid mutations on functions in distant domains.^20,28^

## Supplementary Tables

**Table S1:**
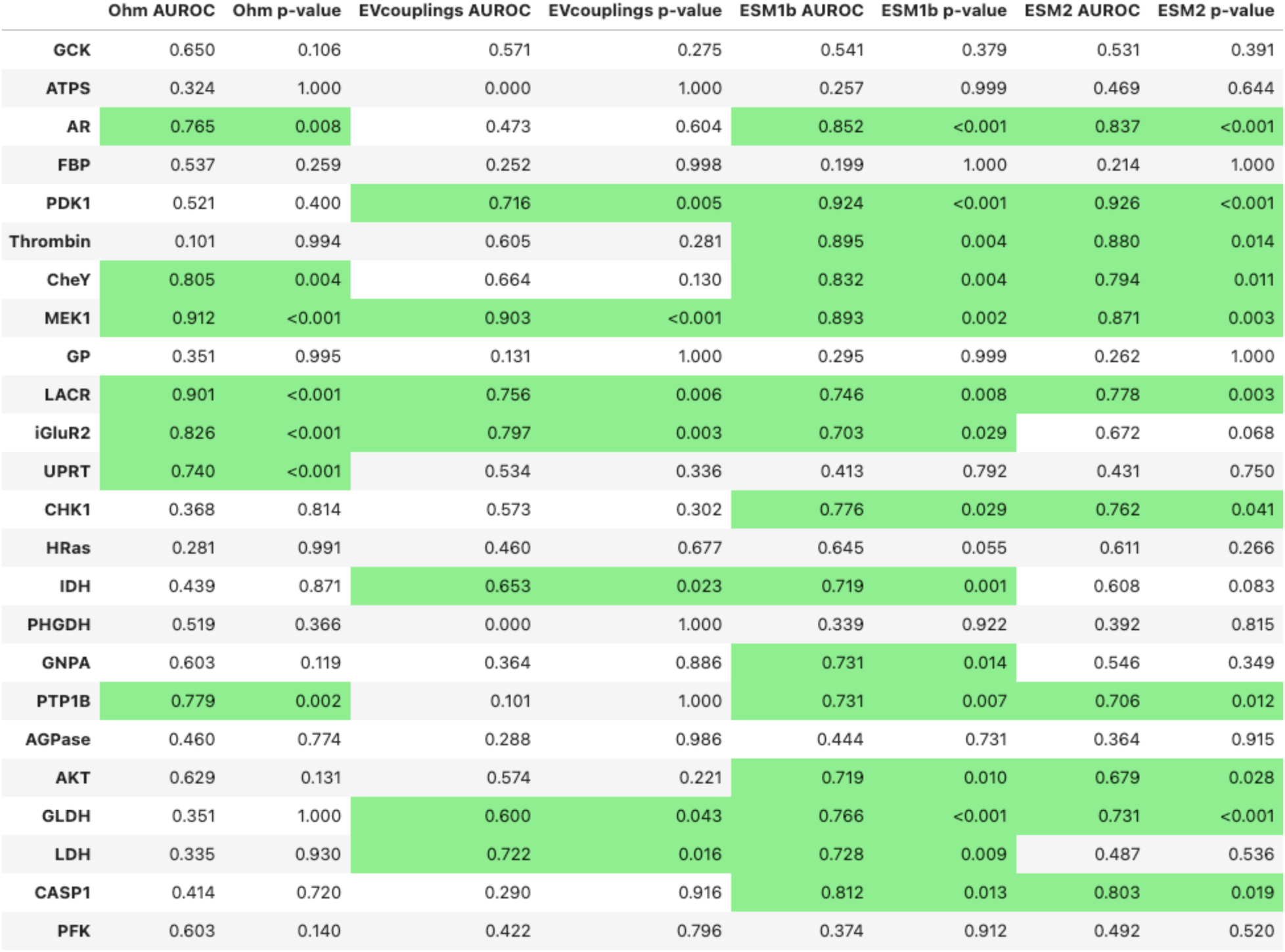
AUROCs and p-values by permutation testing for all proteins for all models. Proteins predicted statistically better than random (p < 0.05) by permutation testing over 1000 permutations are shaded in green (Ohm: 7, EVcouplings: 6, ESM1b: 15, ESM2: 11). Among the 19 proteins from the dataset used in ref. 15: ESM1b identifies 10, Ohm 4, EVcouplings 3 allosteric sites.^15^

**Table S2:** Top 20 highest-attention residues to adrenaline-binding site in B2AR. Residues in B2AR are scored by attention to adrenaline-binding residues, EVcouplings score, and Ohm ACI. Top 20 highest-attention residues to active site are shown here. ‘Motif’ column contains motifs as labeled in ref. 18, and ‘potency’ and ‘efficacy’ columns contain labels indicating changes in function upon alanine scanning in ref. 18.^18^ Rows are colored green if potency or efficacy are reduced, or if residues are part of a known allosteric motif.

| Mutation | GPCRdb numbering | Motif | Potency | Efficacy | ESM1b Attention Rank | EVcouplings Rank | Ohm Rank |
| --- | --- | --- | --- | --- | --- | --- | --- |
| N322A | 7x49 | NPxxY | reduced | reduced | 1 | 184 | 38 |
| D130A | 3x49 | DRY | WT-like | reduced | 2 | 112 | 20 |
| L284A | 6x46 | N/A | WT-like | WT-like | 3 | 117 | 180 |
| W286A | 6x48 | CWxP | N/A | N/A | 4 | 18 | 137 |
| P211A | 5x50 | PIF | N/A | N/A | 5 | 40 | 128 |
| D79A | 2x50 | N/A | reduced | reduced | 6 | 159 | 4 |
| Y219A | 5x58 | N/A | reduced | WT-like | 7 | 165 | 67 |
| W109A | 3x28 | N/A | WT-like | WT-like | 8 | 16 | 197 |
| N318A | 7x45 | N/A | reduced | WT-like | 9 | 56 | 33 |
| C285A | 6x47 | CWxP | reduced | reduced | 10 | 45 | 102 |
| A119G | 3x38 | N/A | WT-like | reduced | 11 | 82 | 159 |
| S319A | 7x46 | N/A | WT-like | WT-like | 12 | 30 | 20 |
| I121A | 3x40 | PIF | reduced | reduced | 13 | 10 | 208 |
| E122A | 3x41 | N/A | WT-like | WT-like | 14 | 13 | 151 |
| Y209A | 5x48 | N/A | reduced | WT-like | 15 | 53 | 159 |
| I205A | 5x45 | N/A | WT-like | WT-like | 16 | 71 | 192 |
| F282A | 6x44 | PIF | reduced | reduced | 17 | 95 | 166 |
| C106A | 3x25 | N/A | N/A | N/A | 18 | 148 | 173 |
| R131A | 3x50 | DRY | WT-like | WT-like | 19 | 213 | 43 |
| Y199A | 5x39 | N/A | reduced | reduced | 20 | 17 | 227 |

**Table S3:**
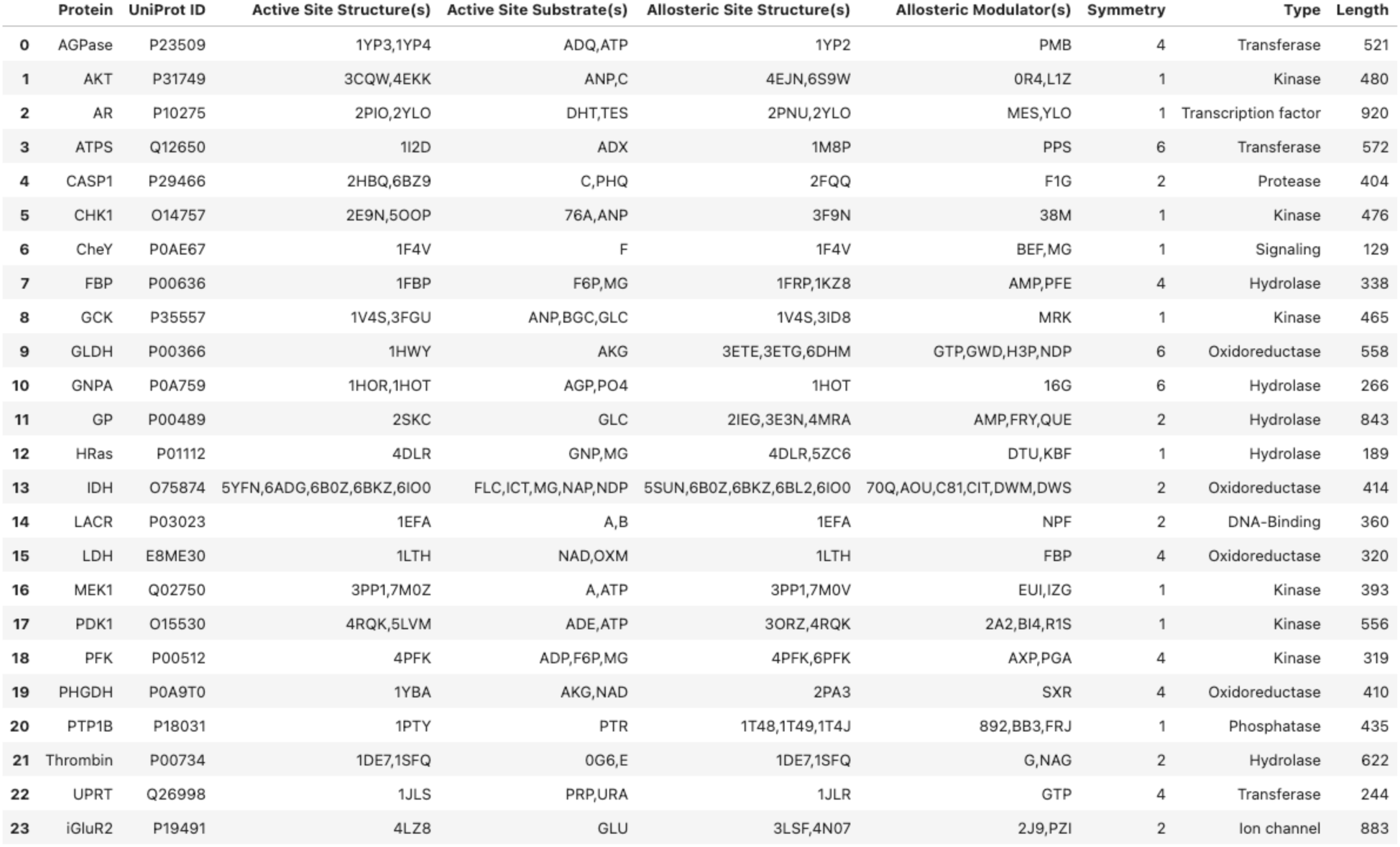
Proteins, structures, and ligands included in benchmark set. Substrate and modulator codes are included as labeled in the corresponding .cif file(s) in the associated structure column. Single-letter codes (i.e. ‘A’) represent protein chains in the structure, while two-letter codes (i.e. ‘MG’) represent ions.

## Supplementary Figures

**Fig. S1:**
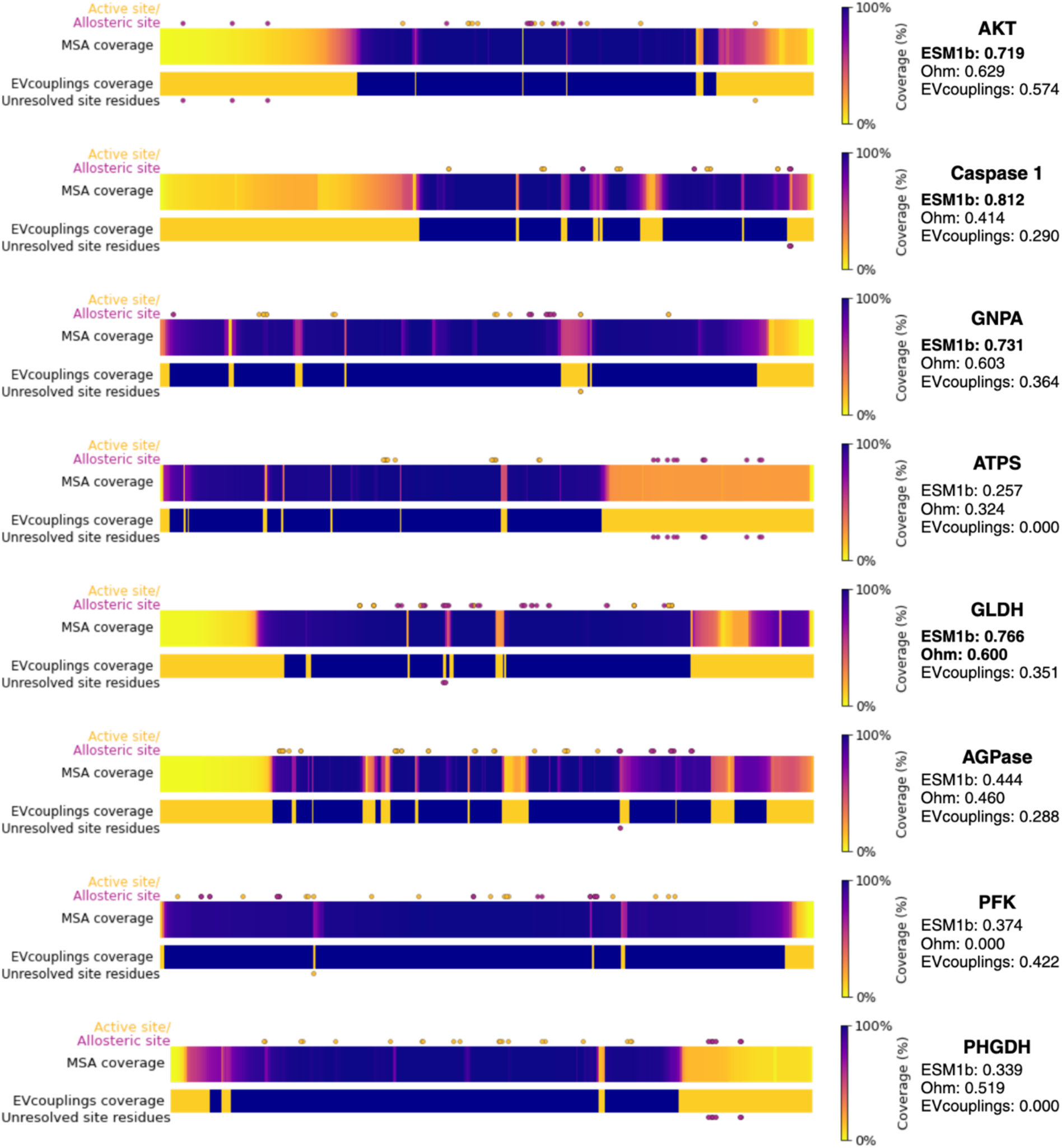
MSA coverage, EVcouplings prediction coverage, and active/allosteric site residues for 8 proteins. For 8 proteins, active site or allosteric site residues are in regions with insufficient MSA depth for EVcouplings to make predictions. Active site (yellow) and allosteric site (purple) residues are shown as dots above each plot, and unresolved residues in MSAs are shown at the bottom of each plot. MSA coverage plots show the percentage of sequences in the EVcouplings-calculated input MSA that contain each residue (top bar for each protein), and EVcouplings coverage plots show positions for which EVcouplings was able to make predictions (bottom bar for each protein). Coverage is represented in binary – the bar is colored blue at positions that EVcouplings covers (can predict at), and yellow at positions that EVcouplings does not cover.

**Fig. S2:**
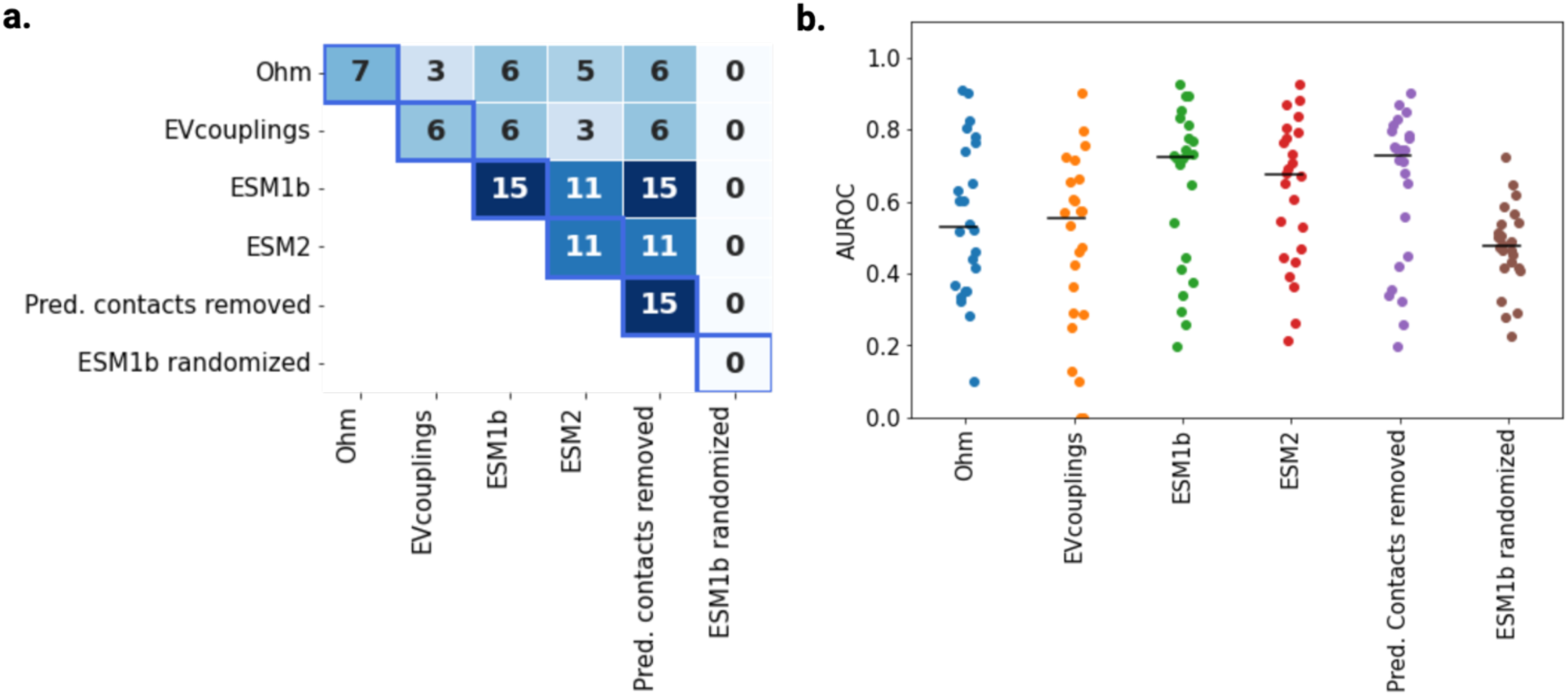
Ablations & controls. (a) Correlation matrix of method performance for ablations. Cells contain the number of proteins (out of 24) predicted better-than-random by each method. Diagonal contains the total number of proteins predicted by each model; for example, ESM1b predicted the allosteric residues in 15 proteins, shown in the (ESM1b, ESM1b) cell. (b) Scatter plot of per-protein AUROCs for each model. Lines depict median AUROC for each model. ESM2 shows worse performance than ESM1b at their largest respective model sizes, despite ESM2 being two orders of magnitude larger by parameter count. This may be due to ESM2 models having been explicitly selected for structure prediction tasks; benefits from scaling may have been somewhat offset by the skewing towards accurate structure prediction, meaning that the high-attention pairs may have been enriched in some layers for contacts instead of allosteric relationships. Performance shows similar performance when the ESM1b-predicted contact map is removed from the attention map pairings before predictions are made. This lack of performance improvement, despite removing some high-attention contacts, may be due to (1) contact maps being represented in a small subset of the layers, and (2) comparatively few of the pairwise positions being affected by contact map removal. Randomly shuffling attention values before predictions are made yields AUROCs roughly equivalent to a random classifier, indicating that the limited post-processing is not driving performance.

**Fig. S3:**
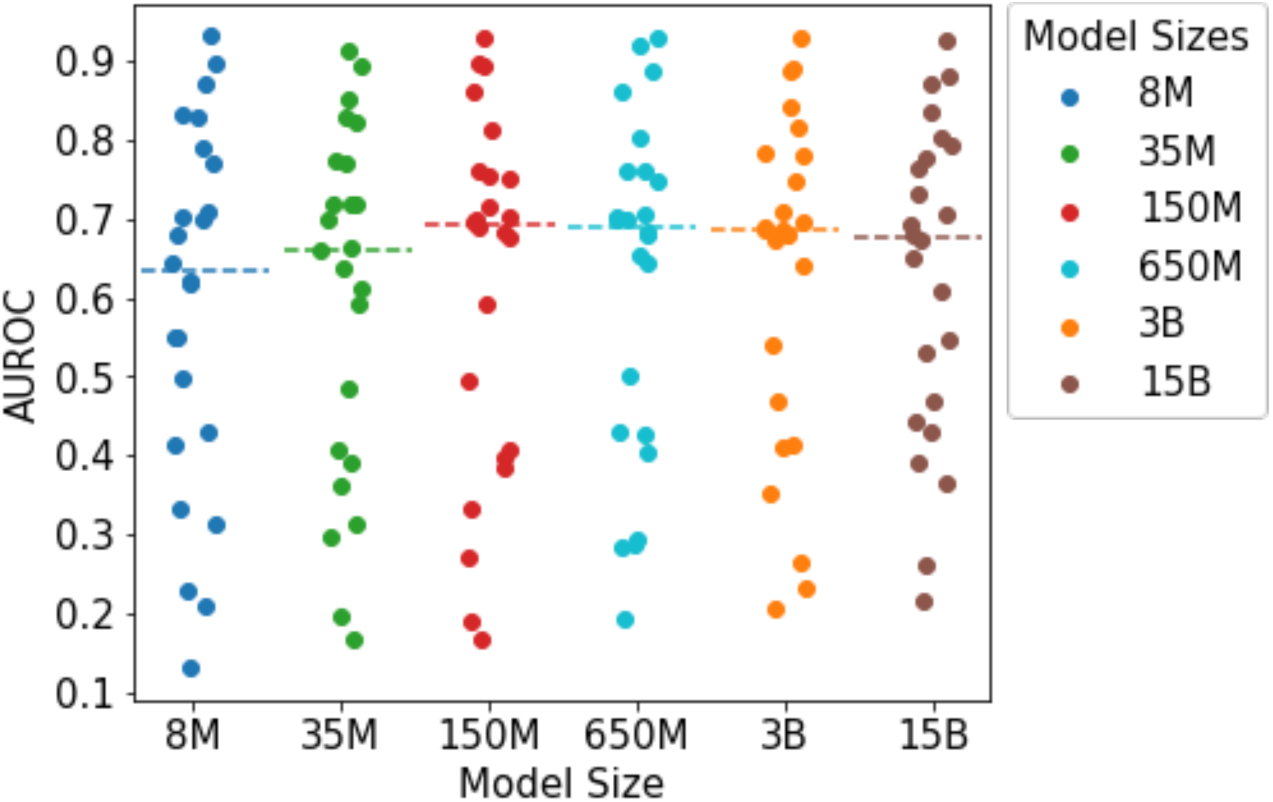
Performance with model scale. Performance improves with size up to ESM2-150M but remains roughly consistent from 150M-15B parameters. There does not appear to be a single model size that drives a large step change in performance. This may be due to ESM2 models being selected for structure prediction tasks.

**Fig. S4:**
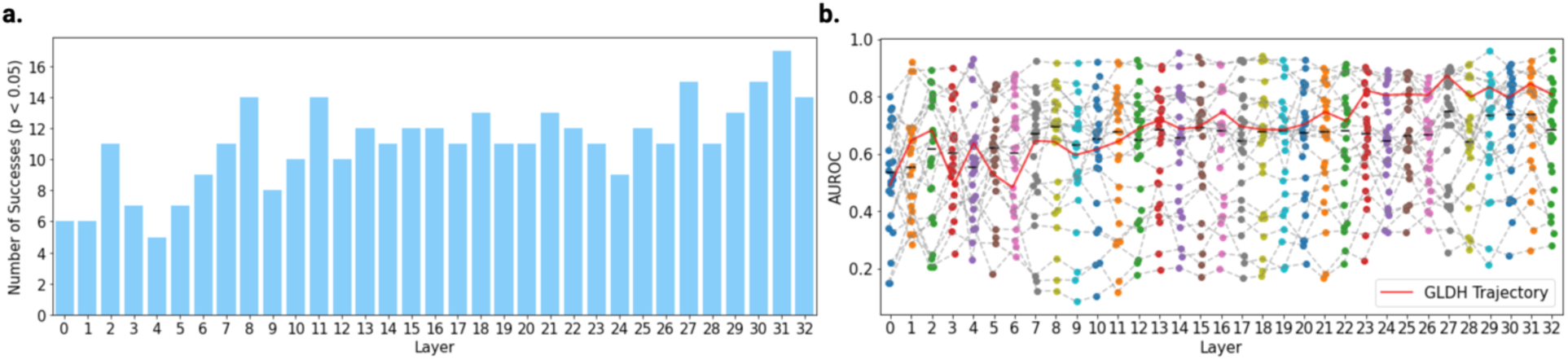
Per-layer performance. Performance of single layers as predictors. (a) Number of successes (proteins with allosteric modulator contact residues predicted better-than-random). Deeper layers appear to identify allostery across more proteins in the benchmark set compared to shallow layers. (b) Each point represents a single protein’s AUROC using the corresponding layer’s mean attention map as a predictor. Generally, predictive power appears to improve at deeper layers. Performance on many proteins is highly variable layer-to-layer. A representative trajectory of predictions on a single protein (Glutamate dehydrogenase) is traced in red.

**Fig. S5:**
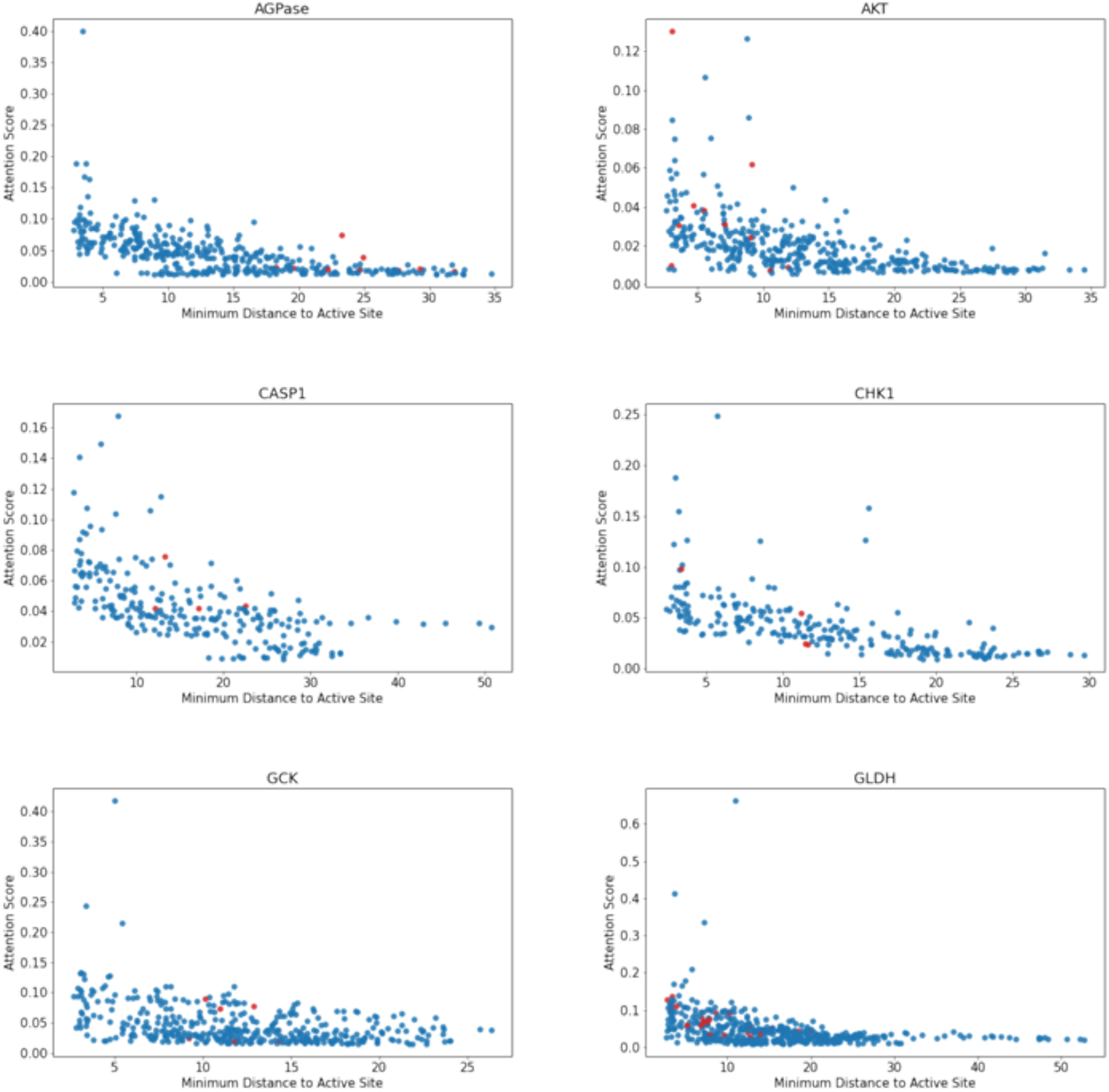
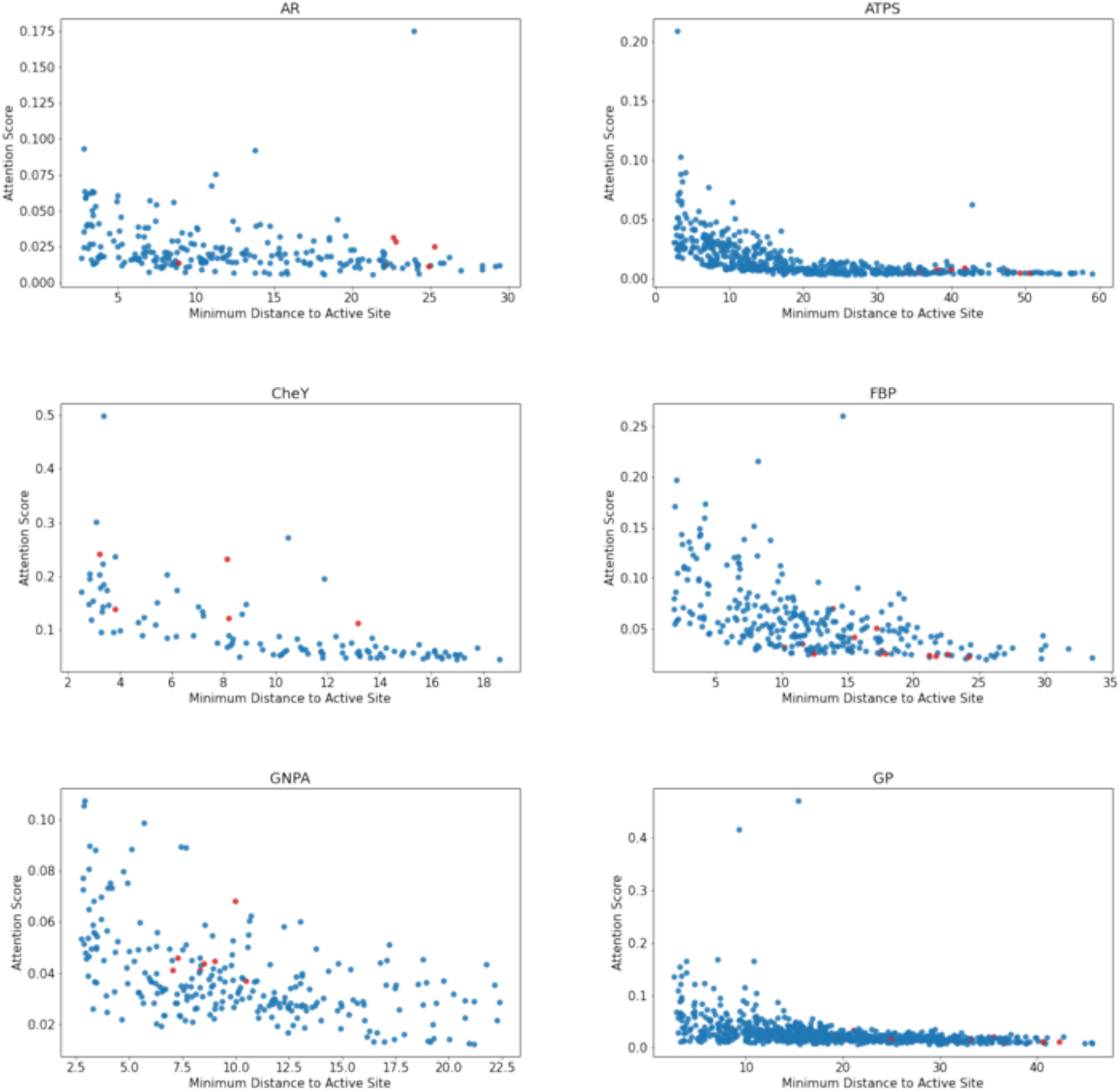
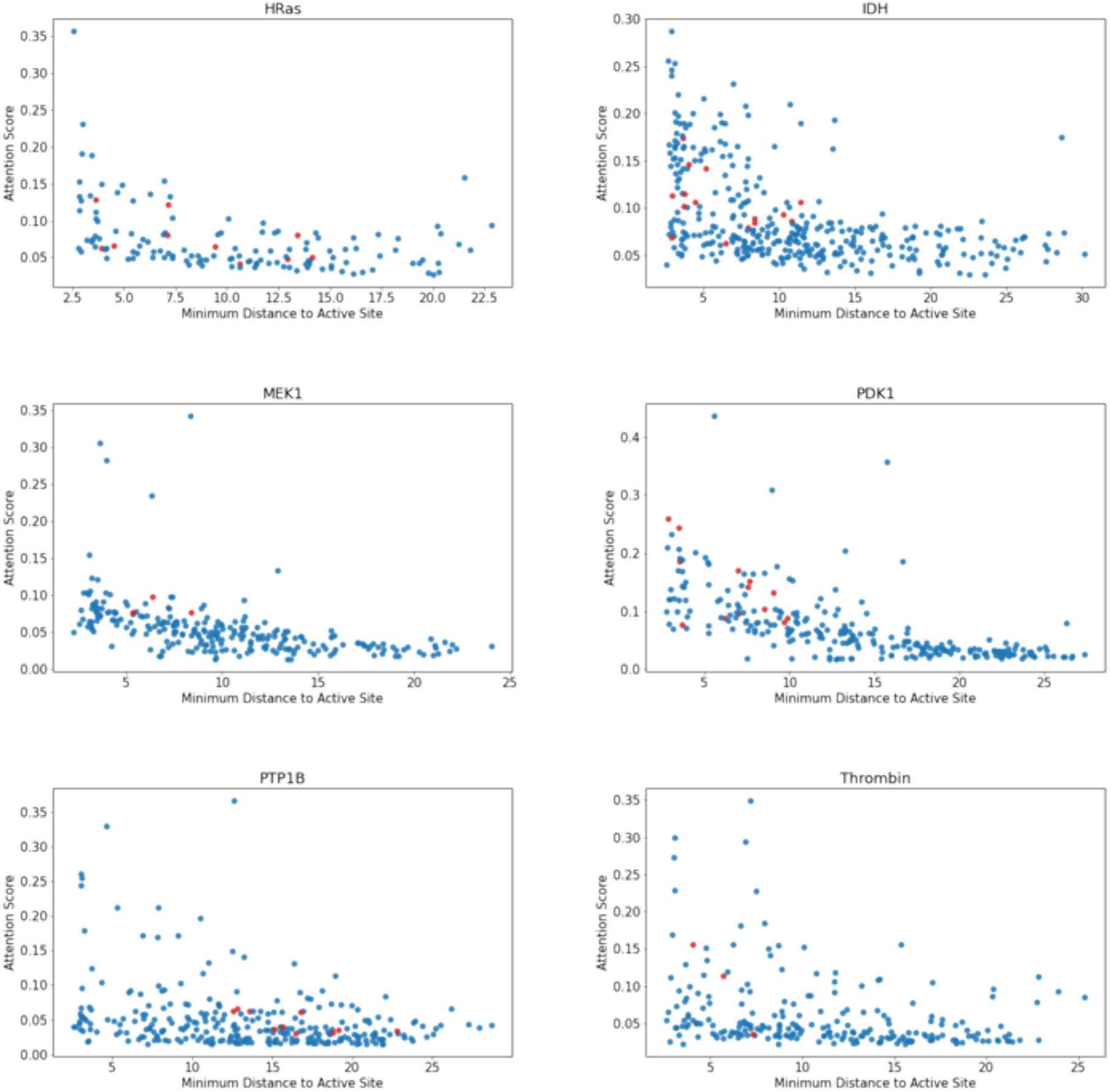
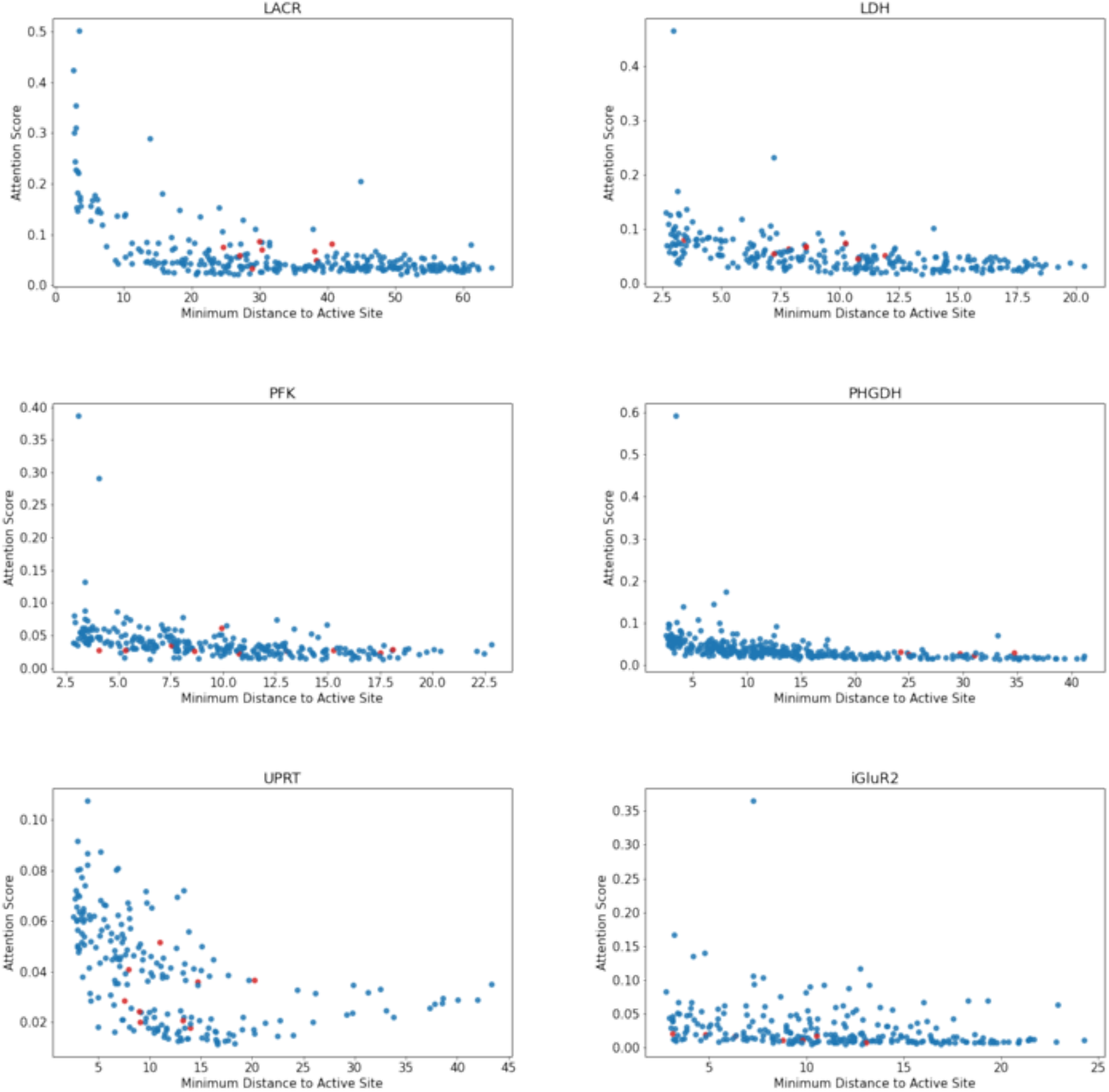
Distance vs. attention score plots for all proteins. Distance from the nearest active site residue (defined as the distance between the closest atom in each residue) on x-axis **(Methods)**. Red dots are known allosteric site residues based on Fig. 2 definition **(Methods)**. Generally, allosteric residues are identified at a range of distances from the active site.

**Fig. S7:**
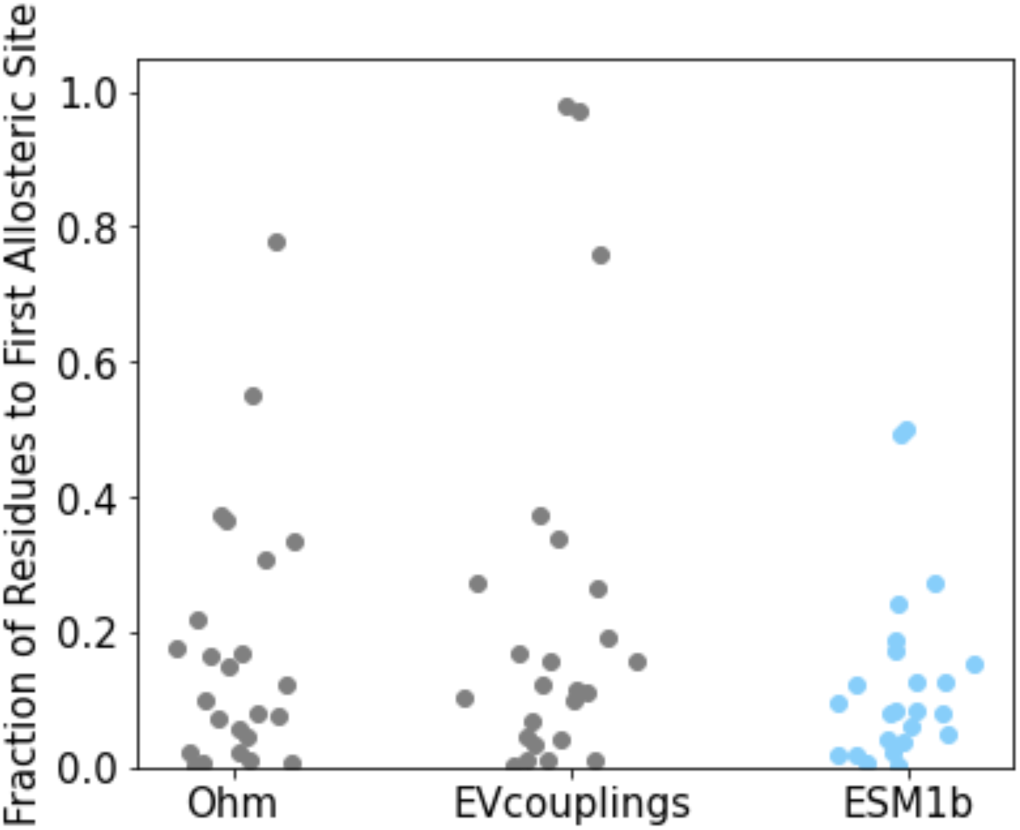
Protein language model ESM1b identifies allosteric sites at low search depth. Depth required in each protein, as a fraction of amino acids in a score-ranked list, to find a single residue in an allosteric site. The protein language model identifies at least one residue in each allosteric site consistently at low depth, performing comparably on this metric to Ohm and significantly better than EVcouplings (p < 0.005).

**Fig. S7:**
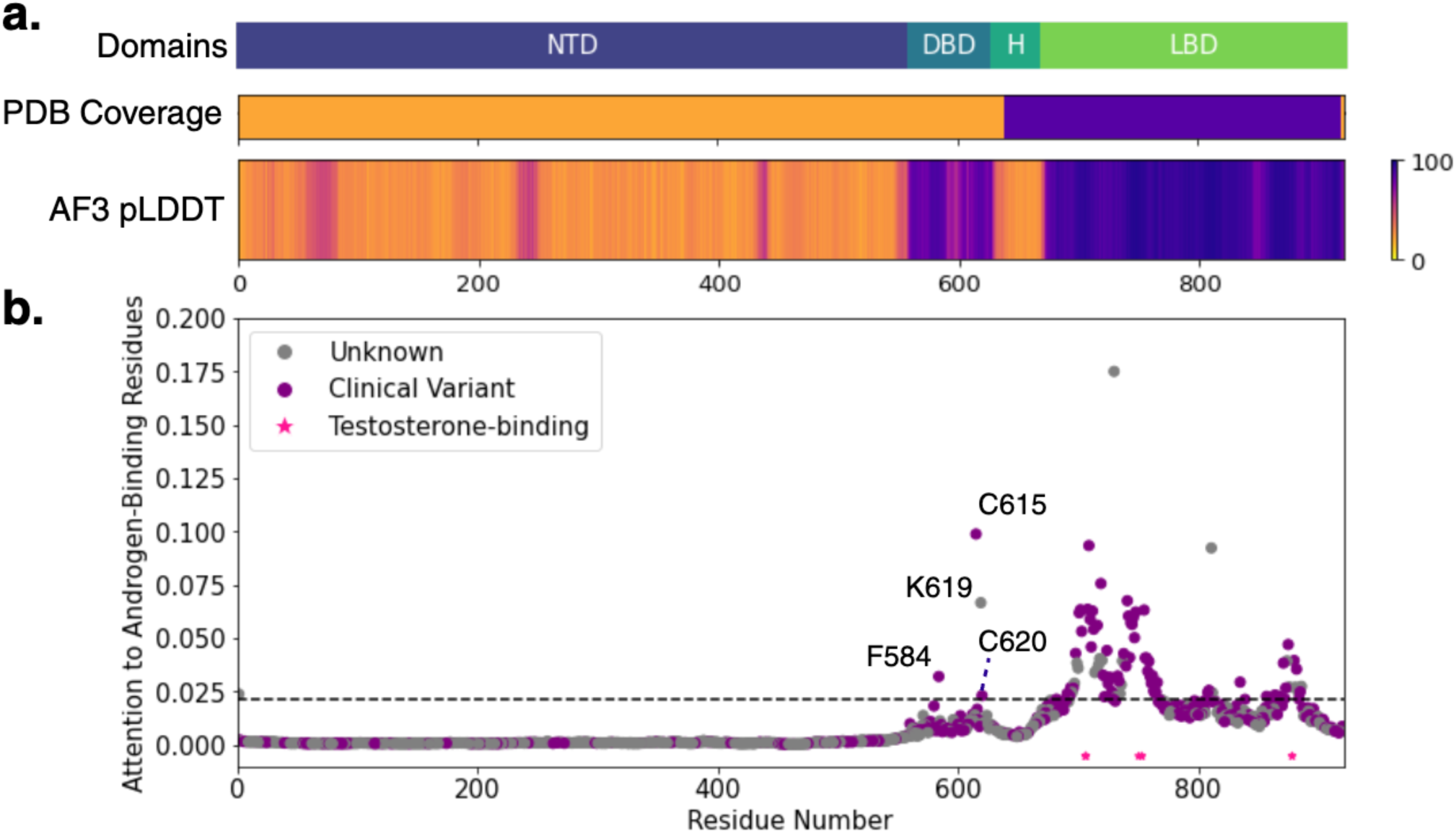
Protein language models identify relationships between distant domains, in the absence of known structure. (a) Top: Labeled domains of Androgen Receptor – NTD: N-terminal domain, DBD: DNA-binding domain, H: Hinge, LBD: Ligand-binding domain. Middle: Crystal structure coverage of human androgen receptor (Uniprot P10275) binarized to 0 (no structures reported) or 1 (present in >=1 structure). There are no reported crystal structures of the DNA-binding domain for human androgen receptor, and no full-length reported structures for any species. Bottom: AlphaFold3 prediction pLDDT confidence score, scaled from 0 (no prediction/very low confidence) to 100 (very high confidence).^54^ (b): Attention of all residues to the testosterone-binding residues in the ligand-binding domain. Disease-associated positions from AndrogenDB are colored in purple (n=347) and unknown positions are colored in gray (n=553).^26^ Pink stars represent testosterone-binding residue locations. High-attention residues (>90^th^ percentile attention score, horizontal line) in the DNA-binding domain are highlighted to explore interdomain communication. C615, K619, F584, and C620 stand out as DBD residues with high attention to the active site; three of these residues are implicated in human disease, and both that have been tested biochemically (C620, F584) were found to diminish androgen binding.^25–28^ In-depth explanations available in supplementary text.

**Fig. S8:**
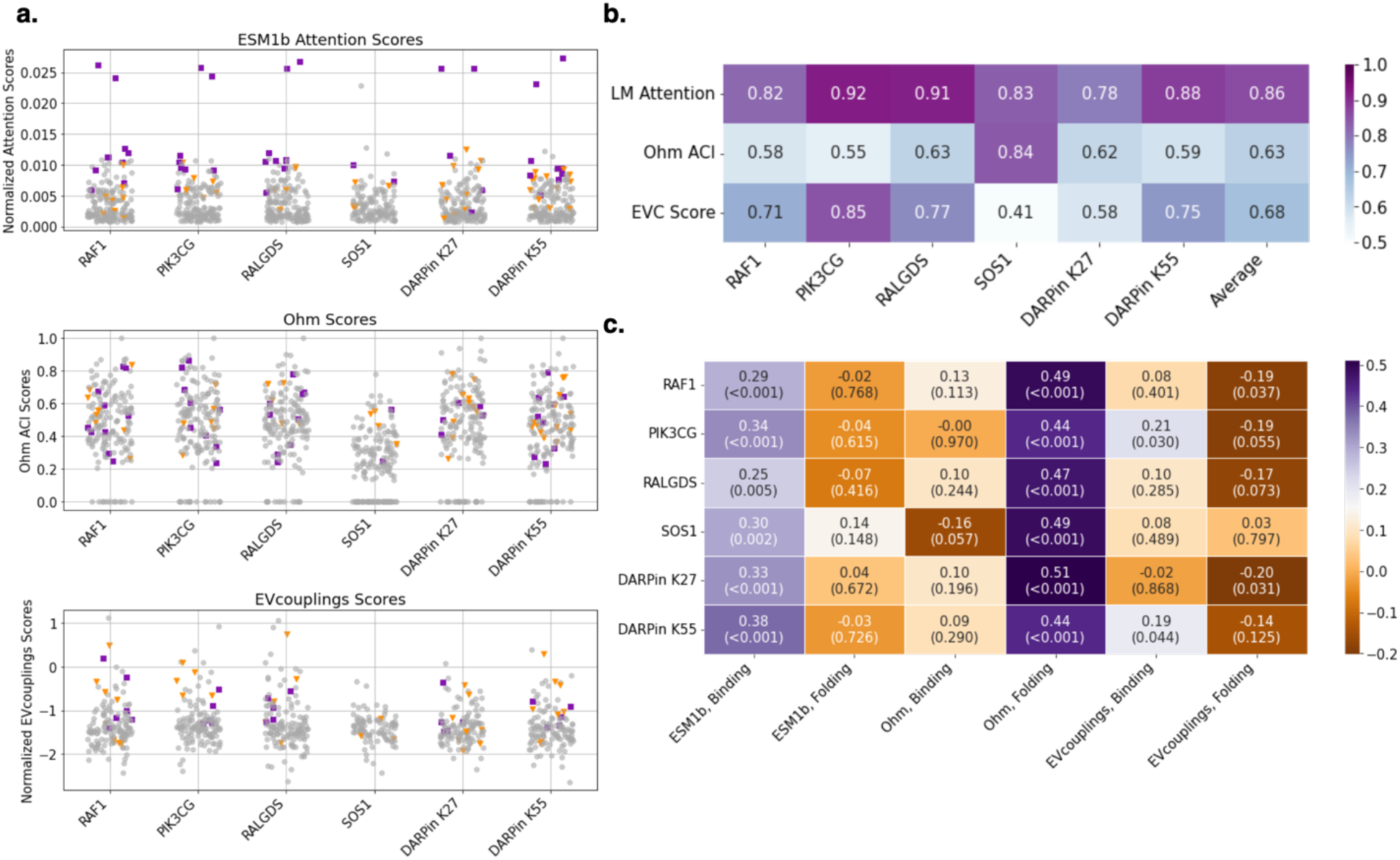
Language models identify allosteric residues in K-Ras accurately and correlate with binding effects, not folding. (A) Top: LM attention-based scores with contact residues to binding partner defined as the active site. Purple squares are GTP-binding site allosteric residues, orange triangles are non-GTP-binding allosteric residues (including some in sotorasib binding site). Binding residues and allosteric residues are defined in Ref. 19^19^ Middle: Ohm scores for each residue plotted for each binding partner. PDB:6VJJ used as input. Bottom: EVcouplings scores for each residue plotted for each binding partner. (B) AUROCs for each model, attempting to classify allosteric residues vs. non-allosteric residues **(see Methods)**. AUROCs are shown for each binding partner, as well as the average AUROC for each model across all 6 binding partners, and scaled from 0.5 (random classifier) to 1 (perfect classifier). (C) Spearman correlations for each model to binding and folding ΔΔG as experimentally determined in Ref. 19^19^ Spearman p-values are represented in parentheses under correlation coefficient *ρ* in each cell. Greater magnitude of *ρ* indicates stronger positive or negative correlation. Language model correlates strongly to binding but not folding, Ohm correlates very strongly to folding but not binding, and EVcouplings correlates weakly in some cases to both.

**Fig. S9:**
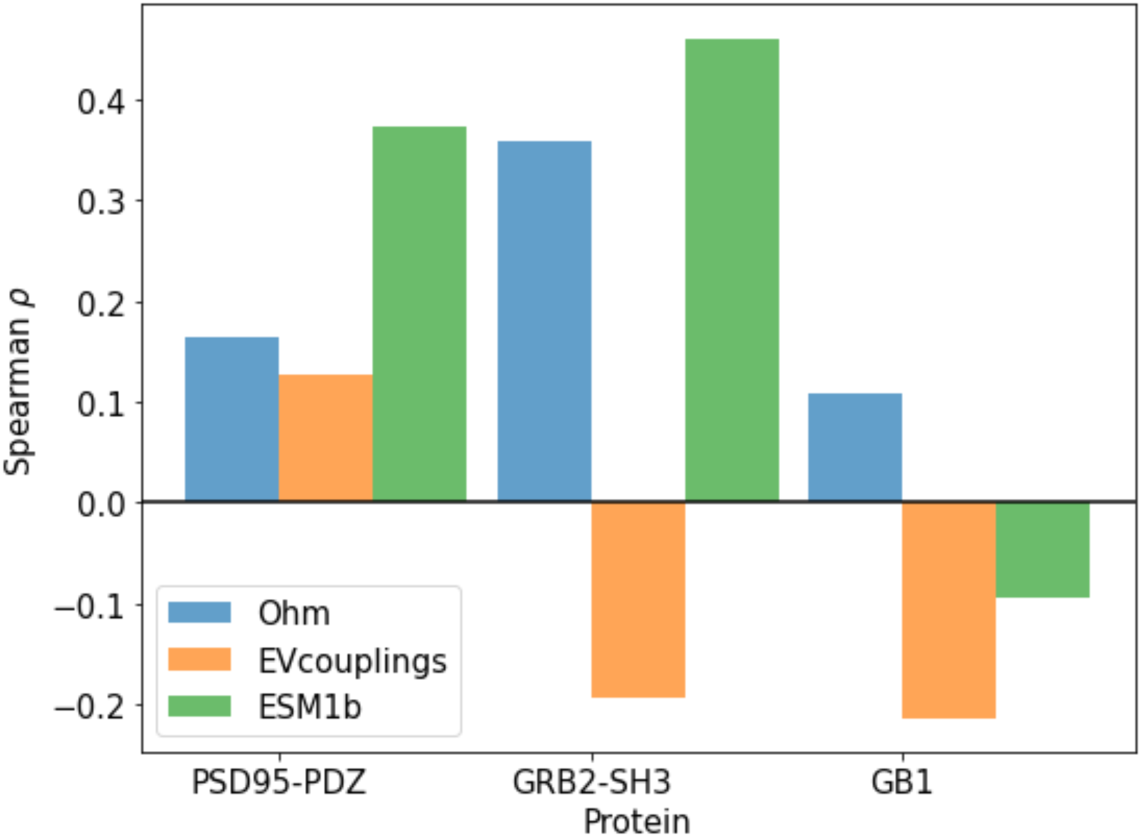
Model performance comparison on deep mutational scanning data. Deep mutational scans probing binding allostery from ref. 29 yield data on allosteric relationships with residue-level resolution for 3 protein binding domains.^29^ We compare model performance for the language model (ESM1b) vs. network model (Ohm) by plotting Spearman correlations of per-residue scores from each model with the number of allosteric mutations at each residue **(Methods)**. Binding residues and allosteric mutations were labeled as in ref. 29.^29^ ESM1b correlates well on PSD95-PDZ and GRB2-SH3, but poorly on GB1. Despite the nature of these examples being small proteins with limited context, and the limited data volume of only having three proteins, these represent a useful validation set as they are fully experimentally labeled as allosteric/non-allosteric at every position.

**Fig. S10:**
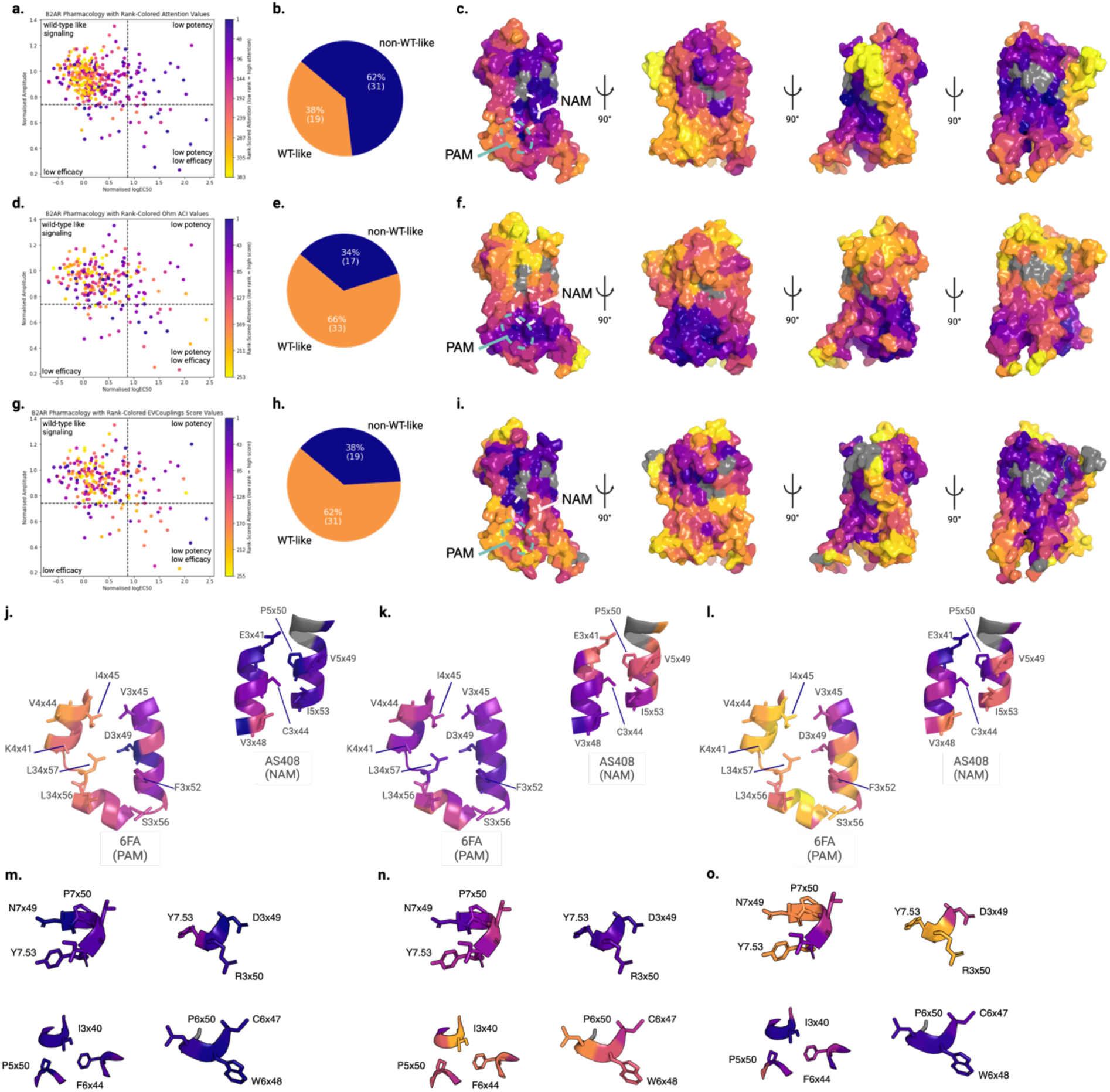
Self-attention identifies residues in all known allosteric sites and motifs in Beta-2 Adrenergic Receptor. Ohm and EVcouplings correlate worse than language model self-attention with experimental mutagenesis data of B2AR. (a, d, g): plot of potency vs. efficacy (data from ref. 18), colored by each model’s rank-scored predictions for each residue (a: ESM1b, d: Ohm, g EVcouplings) among only structurally resolved residues.^18^ Residues in non-WT-like quadrants are ranked higher by language model attention on average. Spearman correlations: (attention, logEC50 = 0.45, attention, amplitude = -0.22). (Ohm score, logEC50 = 0.02, Ohm, amplitude = -0.12). (EVcouplings score, logEC50 = 0.08, EVcouplings, amplitude = -0.04). Spearman p-value < 0.005 for both correlations for the language model, but neither for Ohm or EVcouplings. (b, e, h): Fraction of top 50 residues ranked by attention (b), Ohm score (e), or EVcouplings score (h) in top left quadrant (WT-like phenotype) and other three quadrants (non-WT-like phenotype). (c, f, i): PDB:4LDO colored by each model’s score per-residue (b: ESM1b, d: Ohm, f: EVcouplings). PAM site circled in green, NAM site circled in pink. (j, k, l): PAM, NAM binding sites colored by each model’s score per-residue (j: ESM1b, k: Ohm, l: EVcouplings). Small-molecule contact residues as defined in ref. 18 shown as sticks and labeled by GPCRdb residue numbers.^18^ ESM1b identifies residues in both among the highest-scoring. Ohm identifies PAM, while EVcouplings identifies NAM site. (m,n,o): NPxxY (top left), DRY (top right), PIF (bottom left), CWxP (bottom right) motifs colored by each model’s score per-residue (m: ESM1b, n: Ohm, o: EVcouplings). All four allosteric switches, along with the allosteric sodium binding site (N7x49), are identified as having very high-attention to the adrenaline-binding site by ESM1b.

**Fig. S11:**
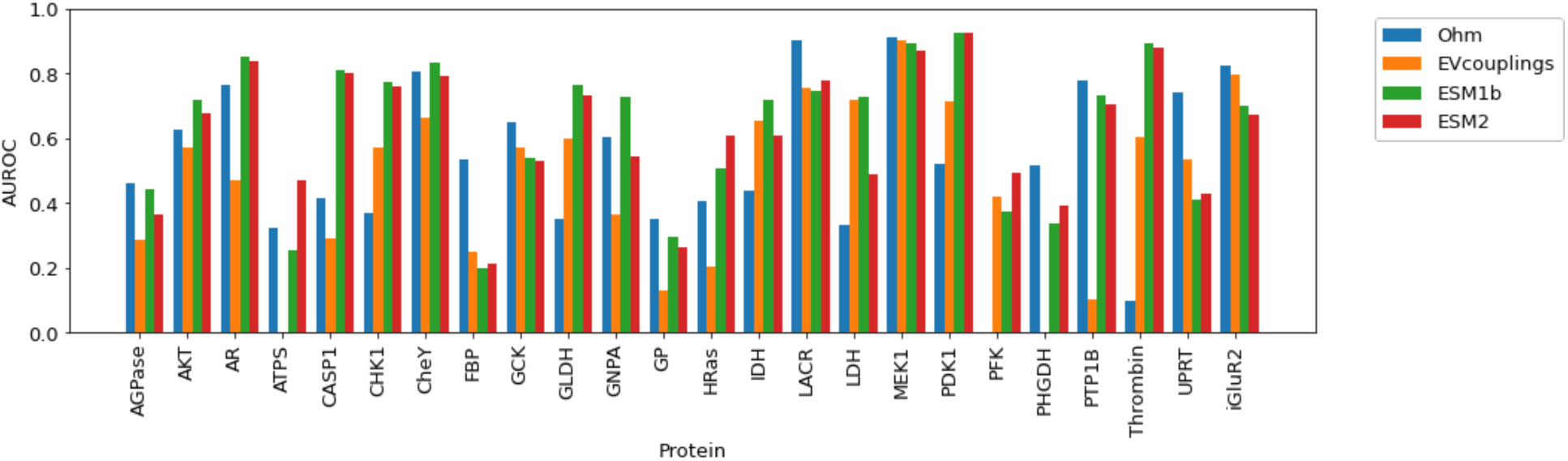
AUROC performance comparison for all proteins in benchmark set. Language models (ESM1b, ESM2) generally outperform the network model (Ohm) and coevolutionary analysis (EVcouplings) on a per-protein basis. ESM1b and ESM2 perform similarly on most proteins.

**Fig. S12:**
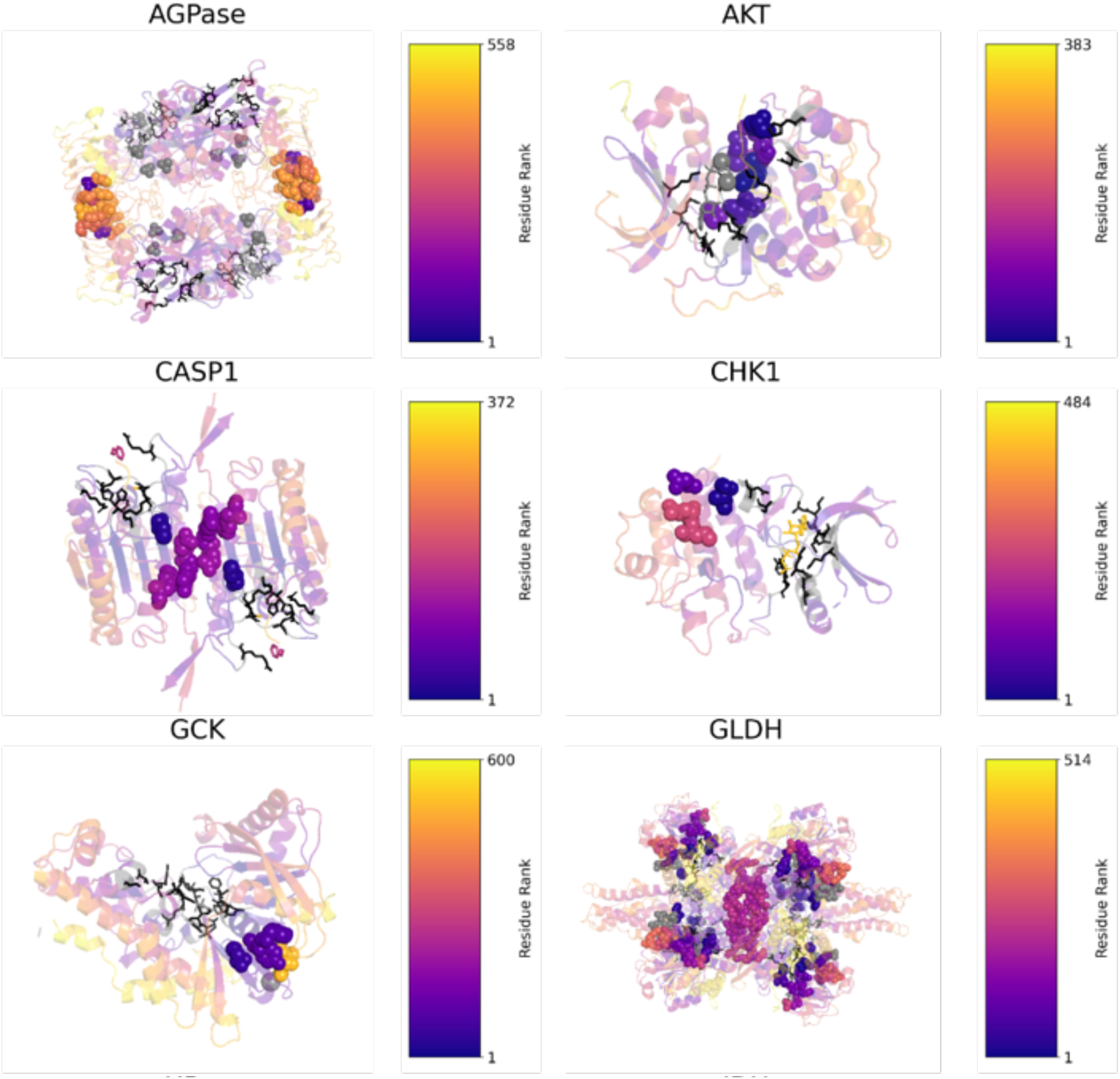
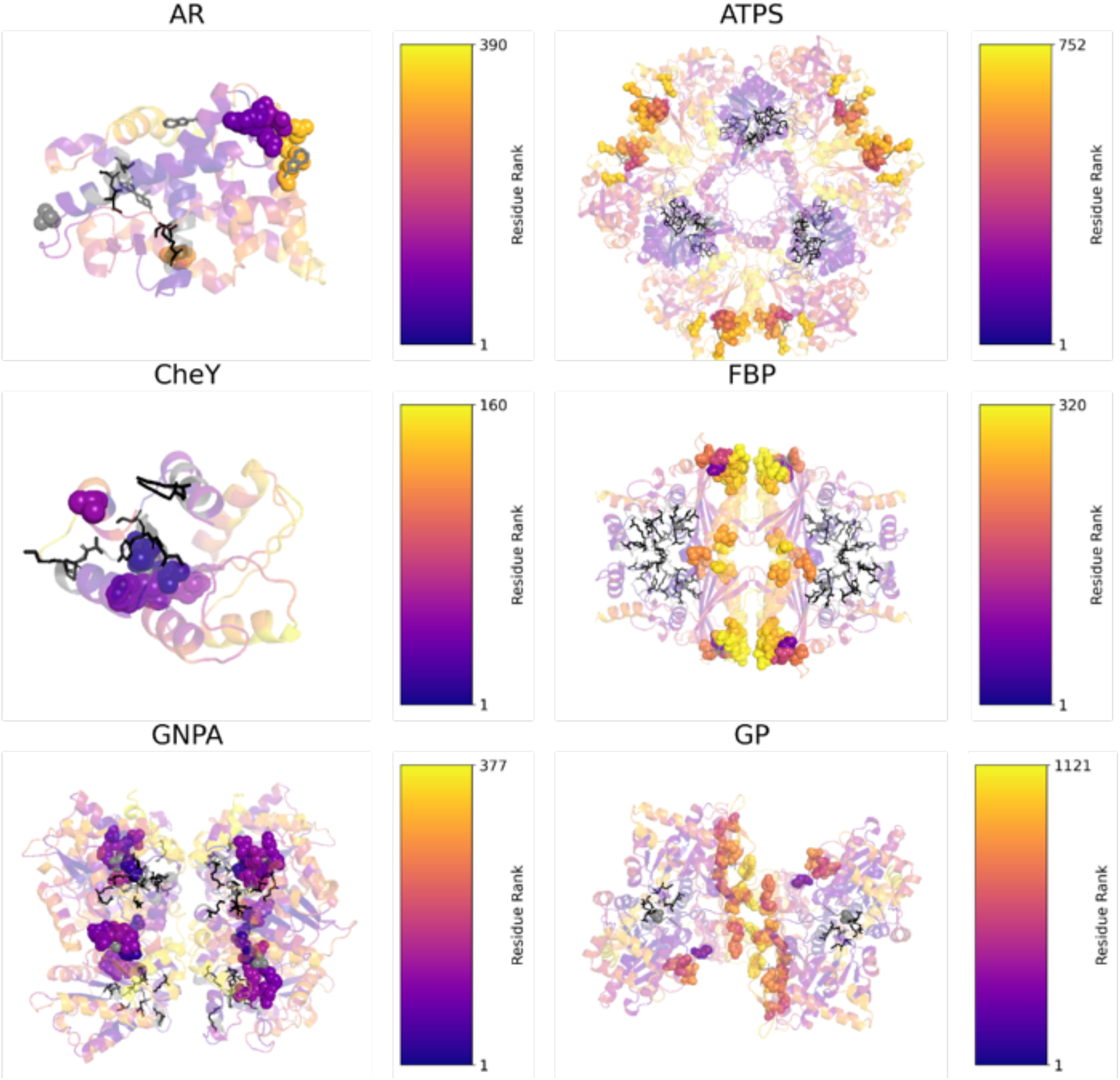
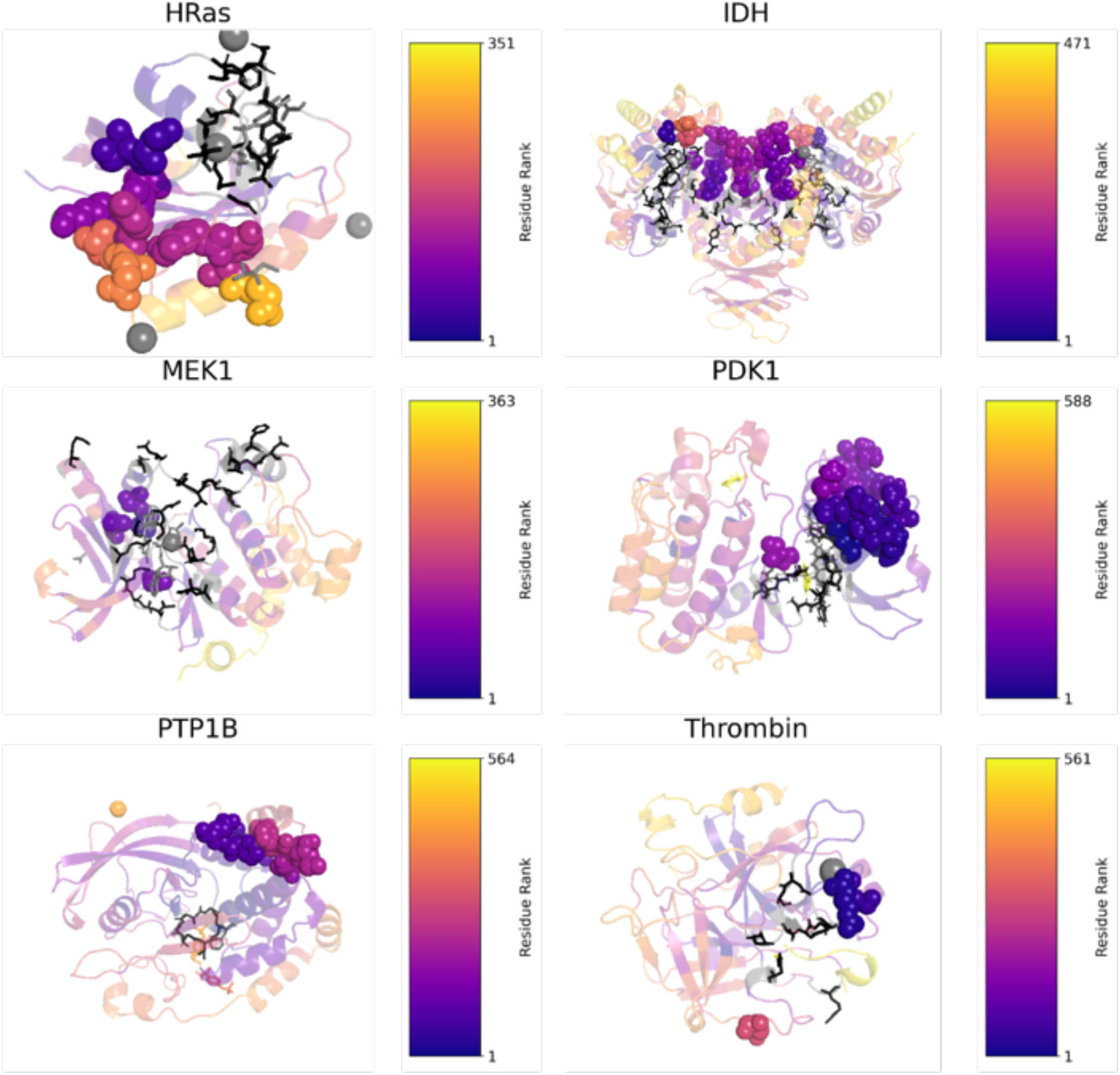
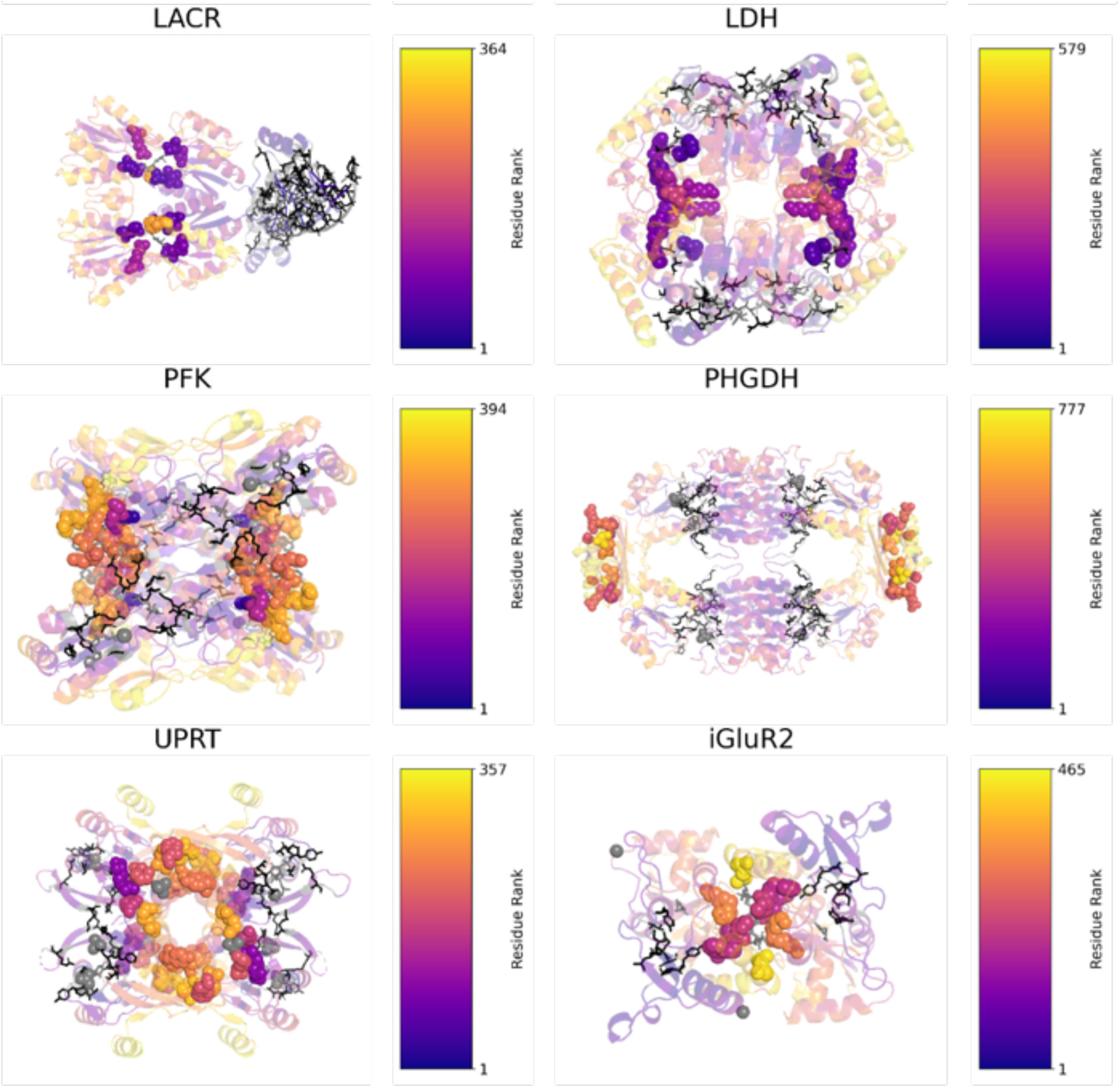
Rank-scored attention coloring for all substrate-bound structures in benchmark set. Crystal structures of all benchmark set proteins with crystallographically resolved residues colored by rank on ribbons (low rank meaning high attention), with purple being highest-attention to active site residues. Allosteric residues (in contact with an allosteric modulator) are represented as spheres and colored. Active site residues (in contact with a substrate) are colored in black and represented as sticks. All other molecules/chains are colored in gray.

